# Charge distribution and helical content tune the binding of septin’s amphipathic helix domain to lipid membranes

**DOI:** 10.1101/2024.07.05.602292

**Authors:** Christopher J. Edelmaier, Stephen J. Klawa, S. Mahsa Mofidi, Qunzhao Wang, Shreeya Bhonge, Ellysa J. D. Vogt, Brandy N. Curtis, Wenzheng Shi, Sonya M. Hanson, Daphne Klotsa, M. Gregory Forest, Amy S. Gladfelter, Ronit Freeman, Ehssan Nazockdast

**Affiliations:** Department of Applied Physical Sciences, The University of North Carolina at Chapel Hill, Chapel Hill, NC 27599; Center for Computational Biology, Flatiron Institute, New York City, NY 10010; Curriculum in Genetics and Molecular Biology, The University of North Carolina at Chapel Hill, Chapel Hill, NC 27599; Department of Biochemistry and Biophysics, The University of North Carolina at Chapel Hill, Chapel Hill, NC 27599; Center for Computational Mathematics, Flatiron Institute, New York City, NY 10010; Department of Mathematics, The University of North Carolina at Chapel Hill, Chapel Hill, NC 27599; Department of Cell Biology, Duke University, Durham, NC 27710; Joint Department of Biomedical Engineering, the University of North Carolina at Chapel Hill, Chapel Hill, NC 27599 and North Carolina State University, Raleigh, NC 27695

**Author notes:** Authors contributed equally (Christopher J. Edelmaier, Stephen J. Klawa, S. Mahsa Mofidi). Correspondence: Experimental, Theory.

## Abstract

Septins are a class of cytoskeletal proteins that preferentially bind to domains of micron-scale curvature on the cell membrane. Studies have shown that amphipathic helix (AH) domains in septin oligomers are essential for septin curvature sensing. Yet, the underlying mechanochemical interactions that modulate this curvature sensing remain ambiguous. Here we use all-atom molecular dynamics alongside a metadynamics enhanced sampling approach to bridge the gap between time and length scales required to optimize and validate experimental design of amphipathic helices. Simulations revealed that the local charge on the termini of an 18-amino-acid AH peptide impacts its helical content and positioning within lipid membranes. These computational observations are confirmed with experiments measuring the binding of synthetic AH constructs with variable helical content and charged termini to lipid vesicles. Taken together, these results identify the helical content of amphipathic helices as a regulator of septin binding affinity to lipid membranes. Additionally, we examined an extended AH sequence including 8 amino acids upstream and downstream of the minimal 18-amino-acid-long AH domain to more closely mimic the native protein in simulations and experiments. Simulations and experiments show that the extended peptide sequence adopts a strong alpha-helical conformation when free in solution, giving rise to a higher affinity to lipid membranes than that of the shorter AH sequence. Together, these results provide insight into how the native septin proteins interact with membranes, and establish general design principles that can guide the interaction of future synthetic materials with lipid membranes in a programmable manner.

**STATEMENT OF SIGNIFICANCE:** Understanding how cells sense and react to their shape is necessary for numerous biological processes. Here we explore the interactions between amphipathic helices, a curvature sensing protein motif, and lipid membranes. Using molecular dynamics simulations, enhanced simulation sampling techniques, and experiments, we find that increasing the helical content of the amphipathic helix or adding charged capping sequences yields higher membrane binding affinity. Understanding these parameters for membrane-binding could enable us to interface and regulate native protein functions, as well as guide the design of synthetic curvature-sensing materials that can interact with and deform lipid membranes.

## INTRODUCTION

Cells have the remarkable ability to both sense their current shape and undergo shape changes as needed to carry out their specific functions. Cellular shape changes are often mediated via interactions of cells with their environment which trigger restructuring of cytoskeletal networks (1). Curvature sensing proteins can promote, stabilize, and transmit geometric information to generate cell responses. Common structural features within these proteins are amphipathic helices (AH domains), which insert into membrane bilayers at lipid-packing defect sites, where there are gaps between lipid headgroups exposing the acyl chains (2). Combinations of experimental and modeling studies support a model in which the hydrophobic portions of AH domains interact with the inner leaflet of membranes. Moreover, it has been demonstrated that a membrane-bound amphipathic helix can maintain a stable folded state (3). While AH domains in various proteins are clearly involved in membrane binding, their binding mechanism and how they directly contribute to other properties such as curvature sensing or higher-order assembly of these proteins are less characterized and not well understood.

Computational studies by the Voth group (4–6) and others (7,8) have been conducted on how AH domains may interact with membranes. One study that focused on the amphipathic helix H0 in endophilin revealed that folding into the helical state is a cooperative process coinciding with the presence of membrane defects on the convex (positive curvature) membrane (4). These simulations show that the folding process is mediated by contacts with lipids that are facilitated by lipid packing defects. Specifically, the helix unfolds when it is not bound to the membrane. The H0 peptide needed to be at the convex membrane surface where more and larger defects are present to fold into an amphipathic helix. In contrast, on flat or concave membranes, the peptide binds the membrane in the unfolded state through hydrophobic interactions and is ‘kinetically trapped’ and unable to form an amphipathic helix. Additionally, it was observed that the defects to which H0 must bind, based on its hydrophobic face when folded, were larger than the defects typically observed for vesicles with a radius of curvature preferred by endophilin. This suggests that as the peptide binds and folds, it may dramatically remodel the defect size and distribution of the membrane.

Other studies have focused on how BAR domain amphipathic helices help organize higher-order structures (5). AH engagement with lipid droplets has also been studied, including simulations of the ‘forward process’ where an unfolded AH-domain in solution folds and binds to a membrane (6). Modeling and experimental work of this nature is not restricted to eukaryotic systems; SpoVM, a bacterial curvature sensor, also contains an AH domain and has been shown to sense/induce membrane curvature (9). As the size and number of membrane defects (defined as exposed hydrophobic lipid tails) seem to play a critical role in protein engagement, computational studies have probed how lipid compositions and curvature define membrane defects which can regulate recruitment of proteins (10,11).

Septins make up a family of conserved proteins, found in many eukaryotic cells, which can sense micron-scale curvatures and direct many other cellular processes (12). In *Saccharomyces cerevisiae*, septin filaments are polymerized from octameric subunits, each of which contains a pair of amphipathic helices (one from each Cdc12 subunit)(13). Given that septin octamers are only 32 nm in size, it has been a puzzle how they sense micron-scale curvatures of the membrane (14). A recent study shows that curvature preference of septins is determined by its multiscale and multi-step assembly (15). Hence, the preferred curvature can change with the available curvatures and protein concentration (15). Cooperative interactions between bound septins and octamers in solution play a key role in the assembly process. This cooperativity could possibly be due to the ability of septins to not only sense but also induce deformations in lipid membranes, allowing additional defect-mediated protein recruitment (16,17). The curvature sensing property of septins has been shown to be mediated via the AH domains as truncating these domains causes septins to lose their selectivity and bind indiscriminately to different membrane curvatures (13). Interestingly, the same studies also showed that expressing two AH domains in tandem is sufficient for conferring curvature sensing properties removed from the context of the native protein. However, the density of bound *single* AH domains increases monotonically with curvature, and does not exhibit an optimal curvature, highlighting the role of multivalent AH domains in sensing specific curvatures and membrane deformation. It remains mysterious how the AH domains of septins promote assembly on such shallow curvatures that are not predicted to produce many or deep membrane defects.

To better understand the curvature sensing properties of these AH domains, we investigate their binding configuration, AH position relative to the membrane, and the helical content of a single AH domain within a planar lipid bilayer. We focus on the septin AH domain from Cdc12 of *Saccharomyces cerevisiae* (Fig. 1A, LEEIQGKVKKLEEQVKSL). Approaching this computationally with atomistic simulations (both on a peptide and full protein scale) is highly challenging considering the mismatch between the membrane binding timescale and the limits of runtimes involved. Based on previous experimental studies, the binding event for a septin octamer occurs roughly every 10-100 seconds in an 11×11 nm square area of membrane (13,15). To atomistically simulate the same size of the membrane area, we can at most achieve a runtime on the order of 100 ns/day. This would result in millions of years to see a single binding event. Coarse-grained simulations with a modest speedup and the ability to sample larger length scales would only reduce the runtime to the order of 10,000 years. To overcome this challenge, we have taken the inverse approach, where we initialize the simulations with a conformation that we hypothesize is nearly in equilibrium, thereby overcoming the energy barriers to binding and reducing the runtime. Based on prior studies, we hypothesize that overcoming the excluded volume effects imposed by lipid head groups is a major energy activation barrier to AH intercalation into the bilayer. To bypass this barrier, a pre-folded peptide is placed at a certain depth within membranes and the corresponding simulations reveal if that configuration is stable. Additionally, we use enhanced sampling methods, namely metadynamics, to find the free energy landscape of the peptide and membrane system to understand where stable equilibrium states could exist.

**Figure 1.**
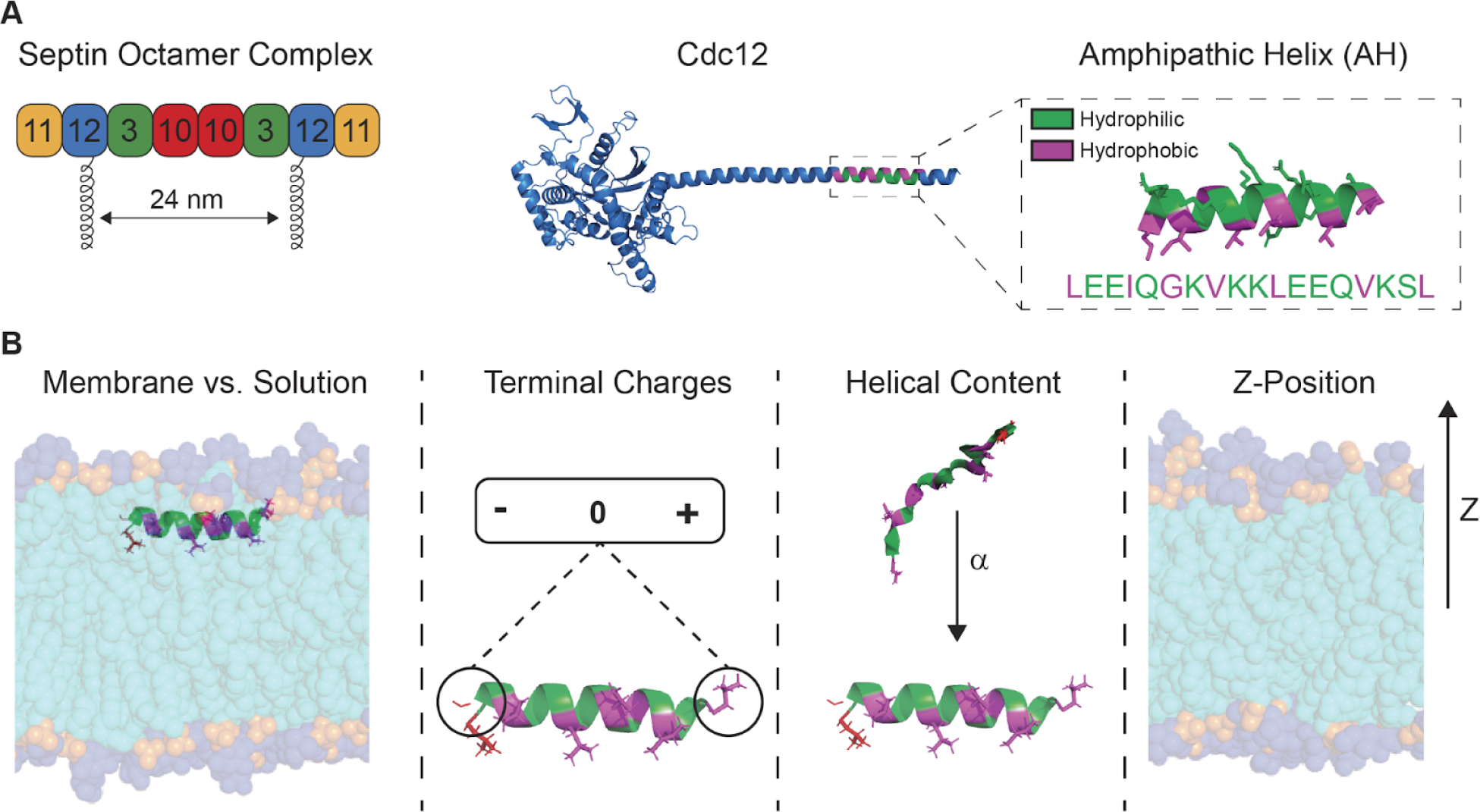
Isolation of Amphipathic Helix and Parameter Space of Binding. **(A)** Septins exist as a palindromic octameric complex that can polymerize end-to-end into filaments. The Cdc12 subunit (AlphaFold model) displays an arm with an amphipathic helix (AH) domain responsible for membrane binding. **(B)** The parameter space explored in our study compares peptides in solution vs. membrane-bound, varying terminal peptide charges, helical content, and Z-position from the center of the membrane.

We varied the local charges on the peptide termini, their helical content, and initial positioning within lipid membranes (Fig. 1B). This allowed us to determine, using molecular dynamics simulations, the quasi-equilibrium states of peptide-membrane binding. Metadynamics is used to determine the free energy landscape of the various peptide-membrane combinations. Experimentally, we synthesized conformationally-constrained AH peptides to pre-define their helical content as a handle to tune structure-dependent membrane affinity. Furthermore, we have tested computationally and experimentally the role of an additional 8 amino acids upstream and downstream of the AH domain on its folding and interaction with membranes. We found that these additional peptide segments at the N and C terminal of the AH sequence have a significant effect in preserving the AH domain helical content in the longer peptide, giving rise to more than a 3-fold higher binding affinity than an isolated AH peptide. These results allow us to design peptides with different affinities for binding membranes *in vitro,* which may enable the construction of materials to programmably bind and deform membranes in the future.

## MATERIALS AND METHODS

### Computational

Simulations were carried out using GROMACS 2022 simulation frameworks (18). All simulations were carried out on the sequence of the budding yeast *S. cerevisiae* Cdc12, the protein that contains the curvature-sensing amphipathic helix (AH) domain (19,20). Initial structures were prepared with CHARMM-GUI using the CHARMM36m force field at a salt concentration of 50 mM KCl and temperature of 300K unless otherwise specified (21–23). The simulations were conducted with a timestep of 2 fs, with Van der Waals (VdW) interactions switched between 1.0 and 1.2 nm, and particle mesh Ewald was used at long distances to compute the electrostatic interactions (24). Hydrogen bonding was calculated via the LINCS algorithm (25). Evolution of the system was carried out using the Nose-Hoover thermostat with a coupling constant (τ*t*) of 1 ps, and a Parrinello-Rahman barostat with a coupling constant of (τ*p*) of 5 ps, reference pressure of 1.0 bar, and a compressibility of 4.5 × 10^−5^. Pressure was coupled semi-isotropically in XY for membrane simulations, and isotropically for peptide-only simulations.

#### Modeling the Amphipathic Helix Domain

Peptide sequences were prepared using PyMOL and GROMACS (18). First, an idealized α helix was prepared in PyMOL based on the 18 amino acid sequence amphipathic helix domain located near the C-terminal of *S. cerevisiae* Cdc 12. Different choices of the capping sequence for the AH domain were used depending on the simulation. Neutral-capped versions had (NH_2_) groups placed at both the N- and C-terminals (CHARMM-GUI membrane builder NNEU and CT2). Charge-capped versions had a positively charged (NH^+3^) group placed at the N-terminal and a negatively charged (O^−^) group placed at the C-terminal (CHARMM-GUI membrane builder NTER and CTER). Note that the net charge of the 18 amino acid sequence for the AH-domain of Cdc12 is predicted to be neutral under our experimental conditions (pH = 7.4).

#### Modeling the Extended AH

We also study the extended AH including 8 residues on either end of the AH domain (34 residues: ERIRLNGDLEEIQGKVKKLEEQVKSLQVKKSHLK). This sequence is located on the C-terminal of *S. cerevisiae* Cdc12. The 18-residue AH domain is in the middle of this peptide with 8 extended residues on both ends. The C-terminal 8-residue sequence on the C-terminus has a +3 net charge while the 8-residue sequence of the N-terminus is neutral under biological conditions (pH∼7). First, we build an idealized α helix of these 34 amino acids in PyMOL and then utilize CHARMM GUI to add neutral cappings for both the N-terminus (NNUE) and the C-terminus (CT2).

Unfolded (melted) AH domains were prepared by simulating the perfect α helical form of the peptide without a lipid bilayer present. This configuration was run through the energy minimization step, followed by equilibration in the NVT ensemble at 300K, with at least 1 nm of solvated TIP3P water surrounding the peptide and 50 mM KCl. The temperature is then ramped up to 500K over 100 ps, run for 10 ns at 500K, then ramped back down to 300K over 100 ps, in order to melt the domain by breaking all of the hydrogen bonds. This final structure is re-equilibrated in the NPT ensemble at 300K for 5 ns to generate a final, unfolded structure.

#### Modeling the Lipid Bilayer

Lipid bilayers were prepared using the membrane builder in CHARMM-GUI (26–28). Each bilayer was composed of a 75:25 mix of DOPC:PLPI lipids, set to be near the size of 11×11 nm^2^ in XY, and solvated with at least 7 nm of TIP3P water above and below the membrane. Systems were relaxed and equilibrated using the CHARMM-GUI-provided procedures (6). Simulations including amphipathic helix domains and extended AHs were initialized in CHARMM-GUI by placing the peptides in the bilayer at different depths. AH-domains are aligned with the bilayer in CHARMM-GUI such that the ILE4 and LEU11 residues are oriented to face the interior, hydrophobic, portion of the bilayer. The same residue alignment was performed for the extended AH to face ILE12 and LEU19 residues downward to the bilayer.

#### Choosing Collective Variables for Analysis

Two natural ways to analyze the system would be the center-of-mass separation of the amphipathic helix and the lipid bilayer, as a measure of excluded volume barrier, and a measure of the helical content of the amphipathic helix. We denote these variables as z and Helical content. The center-of-mass separation distance z is defined as the distance between the middle of the lipid bilayer and the AH-domain carbon-alpha (Cα) atoms. Helical content is measured using PLUMED α*RMSD* collective variable. This is a switching function defined to capture the difference in the distances measured in an idealized alpha helix with the current simulation structure, and represents a number of residues that are thought to be participating in the helical content of the peptide.

#### Enhanced Sampling

Enhanced sampling was accomplished using multi-walker well-tempered metadynamics on our two collective variables z and helical content (29–32). Gromacs coupled with PLUMED was used to simulate these systems (18,33). For each system we ran 4 walkers with an initial Gaussian height of 1 kBT, Gaussian widths of (0.05, 0.1) in CV units, a bias factor of 15, and deposited every 8 ps (4000 timesteps). Neutral-capped metadynamics simulations were started from a configuration with the AH-domain in the surface of the membrane, and charge-capped metadynamics simulations were started from a configuration with the AH-domain in solution. We also included harmonic walls for upper and lower bounds in the z CV to keep the peptide in the simulation cells without moving through the periodic boundary in *z*. These walls were placed at (*z* = 5.0 nm, *z* = −5 nm) with a 1000 kJ/mol/nm force constant (κ = 1000). Free energy surfaces (FES) were constructed in several ways. First, the negative bias was used to reconstruct the FES directly from the hills deposited during metadynamics (34,35). Convergence testing for this first method was done by reconstructing the FES at different percentages of completeness of the metadynamics simulations (75%, 90%, 95%) and comparing the found minima to the final FES (100%). A second method of reconstructing the FES was also used involved in reweighing the original simulation points to find the biased FES, and then using weights to unbias said simulation (34–37). We used both exponential reweighting and Tiwary-based reweighting schemes.

#### Simulations and Analysis

A summary of molecular dynamics simulations can be found in Table S1. A summary of metadynamics simulations can be found in Table S2. Visualizations were carried out using PyMOL and VMD. Simulation trajectories were analyzed in Python using MDAnalysis, MDtraj, and membrane-curvature (19–22). Analysis code can be found online at https://github.com/cedelmaier/dragonfruit.

### Experiments

#### Peptide Synthesis and Purification

Peptide synthesis was performed using an automated Liberty Blue microwave synthesizer with 20% piperidine in DMF as the Fmoc deprotection reagent, and DIC/Oxyma as the coupling reagent. Rink Amide resin was purchased from CEM. Unstapled peptides were made with solid-phase peptide synthesis and amide C-terminal (AH: LEEIQGKVKKLEEQVKSL and 8-AH-8: ERIRLNGDLEEIQGKVKKLEEQVKSLQVKKSHLK). For the stapled peptides, Fmoc-LEE(R8)QGKVKK(S5)EEQVKSL-Resin (0.05 mmol scale) (Fmoc-R8-OH, (R)-N-Fmoc-2-(7-octenyl)Alanine; Fmoc-S5-OH, (S)-N-Fmoc-alpha-(4-pentenyl) alanine, from Ambeed) was synthesized on a Liberty Blue microwave synthesizer. For stapling (olefin metathesis) the procedure is detailed elsewhere (38,39). Briefly, the resin was washed with DMF, DCM and DCE (dichloroethane), then 8 mg 1st generation Grubbs reagent (Benzylidene-bis(tricyclohexylphosphine)-dichlororuthenium) in DCE was added. The reaction vessel was shaken on a shaker for 2 h with slow N_2_ (nitrogen) bubbling. Resin cleavage test indicated that only a trace amount of unstapled peptide was present. The stapling procedure was repeated another time overnight to make sure the reaction was complete. The resin was washed with DCE, DCM and DMF, then Fmoc deprotection was performed. The peptide was cleaved with 95% TFA, 2.5% triisopropylsilane, 2.5% H2O, and purified by HPLC with C18 column and a binary solvent system (solvent A: 0.1% TFA/H2O; solvent B: 0.1% TFA/ACN). Isomers of a and b stapling products were collected separately without specific cis/trans stereochemistry assignment. Final peptide identity and purity was assessed by analytical HPLC (Shimadzu UFLC) and mass spectrometry (Thermo LTQ Ion trap mass spectrometer). Peptides were lyophilized, stored at −20°C and solubilized immediately prior to use.

#### Small Unilamellar Vesicle (SUV) Preparation

All lipids and extrusion accessories were purchased from Avanti Polar Lipids. A lipid solution in chloroform was prepared by mixing the lipid stocks at the appropriate molar ratios (75% DOPC, 25% SoyPI). The lipid solution was then dried under vacuum in a glass vial and resuspended in 16.6 mM HEPES, 50 mM KCl, pH 7.4 by vortexing for 10 minutes (final concentration of lipids is 2 mM). The resulting solution was then extruded with 11 passages through a 100 nm polycarbonate membrane using an Avanti mini-extruder setup. Vesicle sizes were confirmed by dynamic light scattering with a Wyatt DynaPro dynamic light-scattering plate reader.

#### Circular Dichroism

CD spectra were collected on a Chirascan V-100 circular dichroism spectrometer equipped with a temperature control unit (300 K) and using a cuvette with a 1 mm pathlength. Peptides were solubilized in 16.6 mM HEPES, 50 mM KCl, pH 7.4. Peptide solutions were mixed with SUVs (or buffer for samples without vesicles) to yield final concentrations of 50 µM peptide, and 100 µM lipids, and mixtures were incubated at least two hours before collecting the spectra.

#### Fluorescence Anisotropy

Peptides were labeled using BP-Fluor488-NHS ester (BroadPharm) by reacting overnight at a 1:1 molar ratio at 4°C. Free dye was removed by dialysis using Tube-O-Dialyzer systems with a 1K MWCO. SUVs were diluted from the stock to achieve a series of lipid concentrations and subsequently mixed with 25 nM fluorescently labeled peptides for 2 hours. Final buffer conditions were 16.6 mM HEPES, 50 mM KCl, pH 7.4. Fluorescence polarization was measured on a PHERAstar FS microplate reader.

#### Data Analysis and Representation

DLS, CD, and fluorescence anisotropy data were analyzed and plotted using GraphPad Prism 9.4.0. DLS data from each dilution was normalized to create comparable histograms showing size distribution. CD spectra were used as-is with minor smoothing. Fluorescence anisotropy data was normalized to the max polarization for each peptide and the normalized data was fit using the nonlinear fit “specific binding with Hill slope”.

## RESULTS

### Molecular Dynamics Simulations Show Amphipathic Helix Must be Helical to Bind to the Membrane

Previous experiments showed that the AH sequence of septins is necessary and sufficient for membrane curvature sensing but how the sequence engages with the membrane is unclear (13). Here we performed experiments and simulations of the isolated AH sequence to assess how both folding and the biological context of surrounding amino acids impact membrane engagement. A rationale for isolating the AH was to work with a short enough sequence to make atomistic simulations computationally feasible, as well as to explore how certain variations of this short sequence and its folding might influence its binding properties.

When in the context of the native protein, the AH domain is near the C-terminus of the protein but does not have terminal charges from the protein backbone as there are an additional eight amino acids between the AH domain and the C-terminus. However, when isolated from the native context, the charges on the N- and C-termini must be taken into account (Fig. 2A), especially as charges flanking the AH domain can modulate the curvature sensitivity (40). The N-terminus of a peptide contains an amine group which is likely protonated under physiological conditions as the pH is below the average pK_a_ of N-terminal amine groups (41,42). The C-terminus can either be synthesized as a carboxylic acid (negative charge at neutral pH) or as an amide which is neutral. We synthesized a version of the AH peptide (Fig. S1) with a neutral C-terminus to more closely mimic only that domain as it would be in the native protein. Using circular dichroism (CD), we investigated the helical content of the peptide before and after exposure to lipid vesicles (Fig. 2B) which showed that the isolated AH peptide is unfolded in solution (sharp negative cotton effect at 200 nm) and does not change when exposed to vesicles (43).

**Figure 2.**
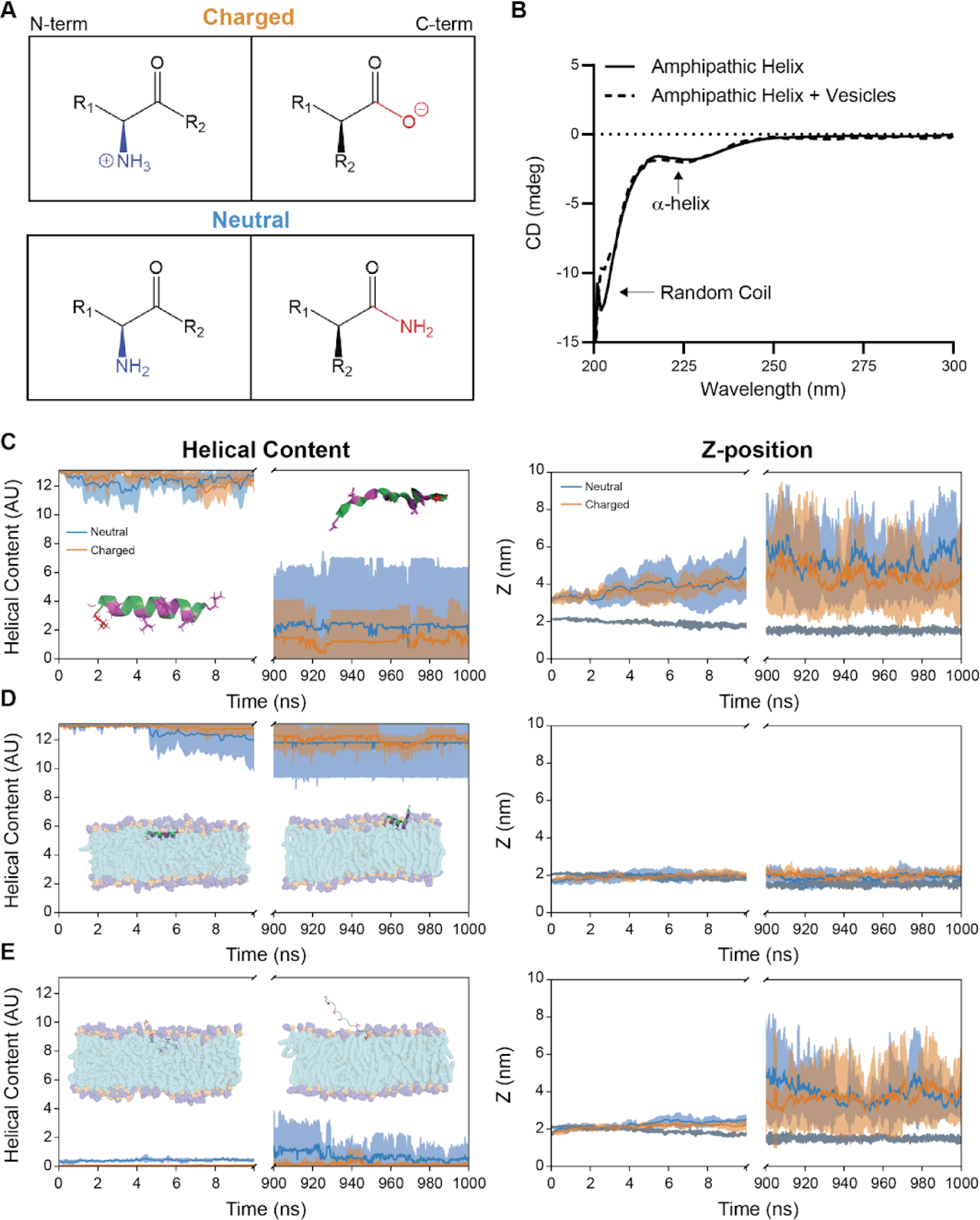
Amphipathic Helix Must be Folded to Bind Lipid Membranes. **(A)** Chemical structures of charged and neutral termini of the peptide. **(B)** Circular dichroism spectra of 50mM peptide with or without 100mM lipid vesicles. **(C)** Trajectories of molecular dynamics simulations showing helical content and Z-position of the peptide when initially folded in solution. **(D)** Trajectories of molecular dynamics simulations showing helical content and Z-position of the peptide when initially folded in the lipid membrane. **(E)** Trajectories of molecular dynamics simulations showing helical content and Z-position of the peptide when initially unfolded in the lipid membrane. Blue represents the neutral peptide, orange represents the charged peptide, gray represents the membrane. Inset snapshots in all cases show the neutral-capped peptide at the beginning and end of a representative simulation.

As an amphipathic helix needs to be folded to display distinct hydrophobic and hydrophilic faces, we hypothesized that the peptide was not binding well under these conditions. To further understand how terminal charges and peptide conformation were important for binding, we explored simulations for neutral and charged peptides with varying initial states: folded in solution above the membrane (Fig. 2C, S2-S3), folded on the surface of the membrane (Fig. 2D, S2-S3), and unfolded placed on the surface of the membrane. (Fig. 2E, S2-S3). The folded peptides that are placed in solution above the membrane were observed to remain in solution and unfold in the timescale of the simulation rather than interact with the membrane. This is consistent with the experimental observations from CD. On the other hand, when the simulations were started with a folded peptide placed on the surface of the membrane, we see both the charged and neutral peptides remain stably bound and maintain a high degree of helical content.

To probe whether helical content is important for stable binding, we initialized the simulation with peptides that are unfolded but placed on the surface of the membrane. In all simulations, the unfolded peptides were ejected from the membrane into solution and remained unfolded. Together this suggests that in the native context, the AH domain binds to the membrane by folding into a helical conformation and matching their hydrophobic and hydrophilic domains with that of the lipid bilayer, but this state is not achieved when the peptide is isolated from the protein since it unfolds.

### Enhanced Sampling of Free Energy Landscapes are Validated Experimentally

To quantify different possible AH interactions with the membrane, we measured the energy landscape of the AH peptide binding to a lipid membrane in simulations with respect to Z-position and helical content. We carried out multiple walker well-tempered metadynamics simulations of the AH peptides with neutral and charged caps to calculate a free energy surface (FES) (Fig. 3A,B, S4-S5). The membrane surface is still found roughly at 2 nm in Z. Free energy differences are determined with respect to the minimum value shown in the plot, which does not include the lowest α bin, and we only plot the first 100 kJ/mol to highlight changes that would otherwise be overshadowed by the very energetically unfavorable states. The lowest α bin is excluded because it includes edge effects due to the switching function used to calculate alpha.

**Figure 3.**
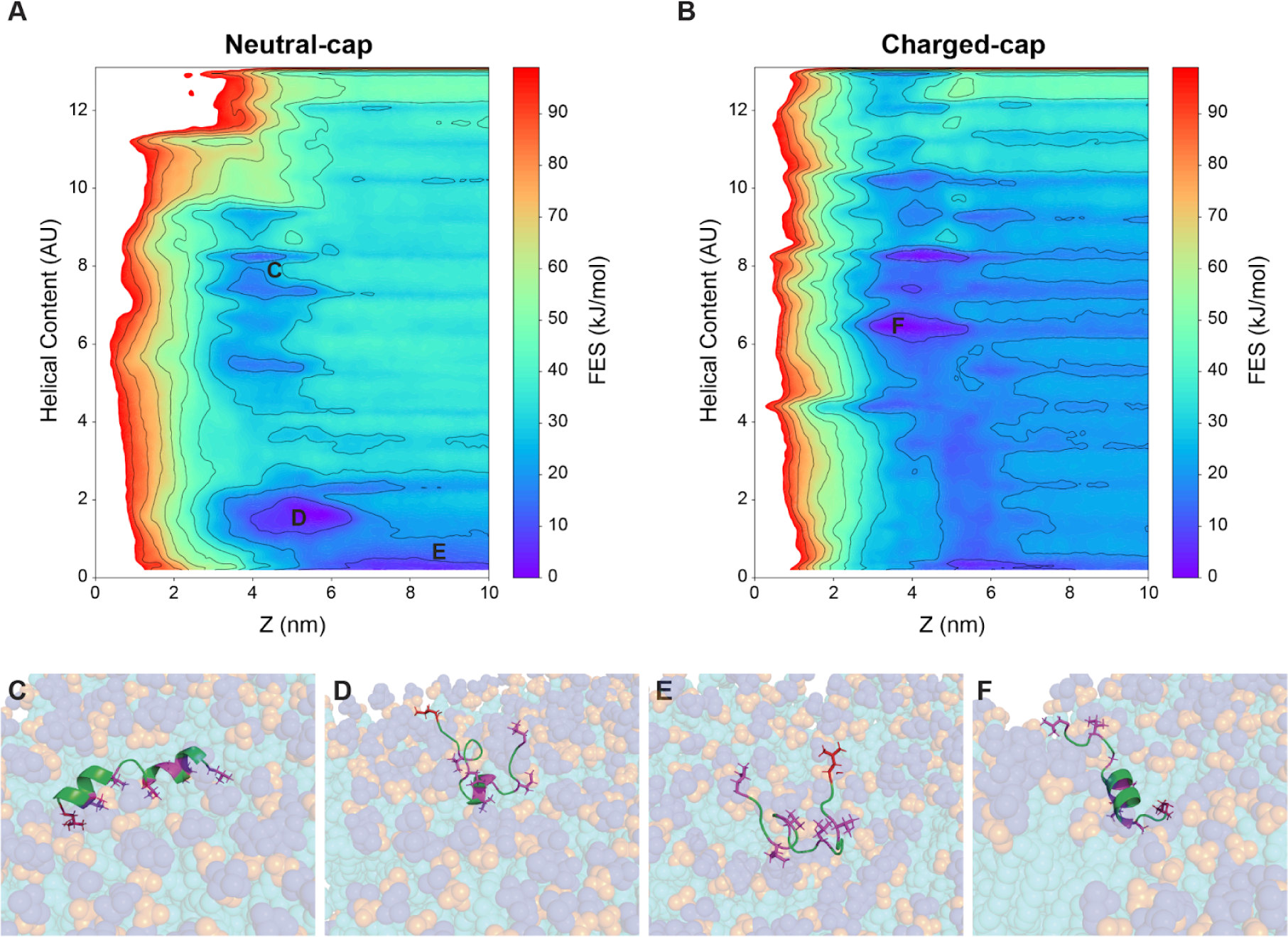
Free Energy Surfaces From Metadynamics. **(A)** Free energy surface reconstruction of the neutral amphipathic helix. **(B)** Free energy surface reconstruction of the charge-capped amphipathic helix. The surface of the membrane is approximately at z=2 nm. **(C)** Metadynamics simulation snapshot of neutral capped peptide at Z=2.1 nm, helical content 8.3. **(D)** Metadynamics simulation snapshot of neutral capped peptide at Z=2.7 nm, helical content 1.7. **(E)** Metadynamics simulation snapshot of neutral capped peptide at z=4.5 nm, helical content 0.5. **(F)** Metadynamics simulations snapshot of charge-capped peptide at Z=1.7 nm, helical content 6.4.

For the neutral-capped AH domain (Fig. 3A), several minima appear. We see a minimum in the membrane at a relatively high helical value (Helical content ∼8) (Fig. 3C) which is in agreement with the expected stable state from the MD simulations. There is also a minimum near the membrane but with low helical content (Fig. 3D) which is likely mediated by interactions other than the hydrophobic insertion of one side of the AH domain. Lastly, we also observe a minimum in the FES that is far from the membrane and a low helical state (Fig. 3E) corresponding to the MD results of the peptide started in solution. We can calculate an approximate ΔG from different minima as well. The FES difference between the ‘free’ form of the peptide (Fig. 3E) and the minima near the membrane, but not at high helical content (Fig. 3D) is ΔG_ED_ = −12 kJ/mol in favor of the ‘near membrane’ state. However, the energy difference to the ‘helical inserted’ form (Fig. 3C) of the peptide is unfavorable from both of these other minima (ΔG_EC_ = 3.5 kJ/mol and ΔG_DC_ = 15.5 kJ/mol). This calculation is in agreement with the AH binding being a very slow and improbable event.

Charged-capped peptides (Fig. 3B) lead to large differences in the energy map of the AH-domain binding to the membrane, as can be seen by comparing the structure of the FES. The location of the membrane has not significantly changed, and is still found at roughly 2 nm in Z. The main difference observed with the charged-capped peptide FES is that the helical states in solution are more accessible as compared to the neutral capped peptide for many values of helical content, which can be seen by comparing the energy gradients in solution along the helical content axis. Unlike the neutral-capped AH-domain, where the minimum FES was observed in unfolded state and in solution and near the membrane (see Figure 3A), a global minimum now appears around (Z ∼ 2 nm, α ∼ 6) corresponding to a surface bound peptide in a half-folded configuration (Figure 3F). Using the same definition of ‘solution’ as the neutral-capped peptide (Z ∼ 4.5 nm, α ∼ 0.5), we can calculate a ΔG = −16.6 kJ/mol in favor of the membrane-bound and helical configuration. The FES results show a very large preference to membrane binding in charge-capped AH domains, compared to the neutral case.

Note that there is no energetically favorable state in solution for the charge-capped peptide, only one near the membrane at low helical content. However, the energy barriers between these states are of order kT and thus accessible through thermal fluctuations. In both charged and neutral-capped peptides, it is very energetically unfavorable for the peptide to be buried in the hydrophobic region of the membrane, as can be seen by the steep gradient in Z < 2 nm. Taken as a whole, this FES suggests that a charge-capped AH-domain would fluctuate between the unfolded form in solution and an associated state at the membrane surface more easily than the neutral capped peptide.

To confirm that the metadynamics had sufficiently converged, we performed two tests. First, changes in the FES were observed at 75%, 90%, 95%, and 100% complete, and the minima were compared (Fig. S3) (44). Two minima can be seen developing and remaining roughly stable depending on the percentage completeness of the simulations. This gives us confidence that the metadynamics algorithm has converged. Additionally, we constructed the biased free energy surface directly from the multiple walker trajectories (Fig. S4). The flatness of this surface gives us further confidence in the convergence of the algorithm. Surface variations can be roughly interpreted as the error in the free energy calculation. Unbiased free energies can be constructed by going through a reweighting procedure. Exponential and Tiwary reweighting gave the same FES (30,34,35,37). These reconstructed surfaces generate a FES that is quantitatively very similar to the original FES (generated by summing up the Gaussian hills deposited during the metadynamics algorithm).

We also carried out simulations of the system in traditional MD, starting from bound configurations found on the neutral-capped free energy landscape (Figure S6, X1−X4), but without the underlying bias potential from metadynamics. Short 300 ns simulations confirmed that the peptides converged to the local free energy minima (or at least traveled along the local gradient of the free energy surface); see the initial and final states of the AH domains in Figure S6. We observe that for X1, which corresponds to a bound unfolded AH (Z ∼ 1.9 nm, α ∼ 0) unbinds and ejects to the solution (z ∼ 6.0 nm, α = 0.6). Similarly, X2 (Z ∼ 1.9 nm, α ∼ 4) transitions across the local free energy basin and folds further by the end of the simulation (Z = 1.6 nm, α ∼ 8.7). X3 (Z ∼ 1.9 nm, α ∼ 8) remains roughly stable, ending at (Z ∼ 2.0 nm, α ∼ 8.5). X4 (Z ∼ 1.9 nm, α ∼ 12) starts outside of the displayed FES (energetically an unfavorable state), but transitions into the shown space ending at (Z ∼ 2.0 nm, α ∼ 11.2). As expected, the low helical content simulation unbinds from the membrane region, while higher helical content simulations transition towards the FES minima found near the surface of the membrane.

To test predictions from these simulations we designed an experimental system that would allow us to modulate helical content of the AH domain and measure membrane binding. To do this we synthesized the AH peptide with a charged N-terminal (amine will be protonated at neutral pH), a neutral C-terminal and incorporated a hydrocarbon staple to regulate helical folding by mutating the 4th and 11th amino acids (i, i+7 spacing). The modified amino acids are covalently coupled together to form a staple forcing a helical conformation (Fig. 4A, S7) (45). As the staple contains a carbon-carbon double bond, this will result in two different isomers (cis and trans) with varying degrees of helical content which will allow us to correlate helical content with the ability of the peptide to bind a lipid membrane.

**Figure 4.**
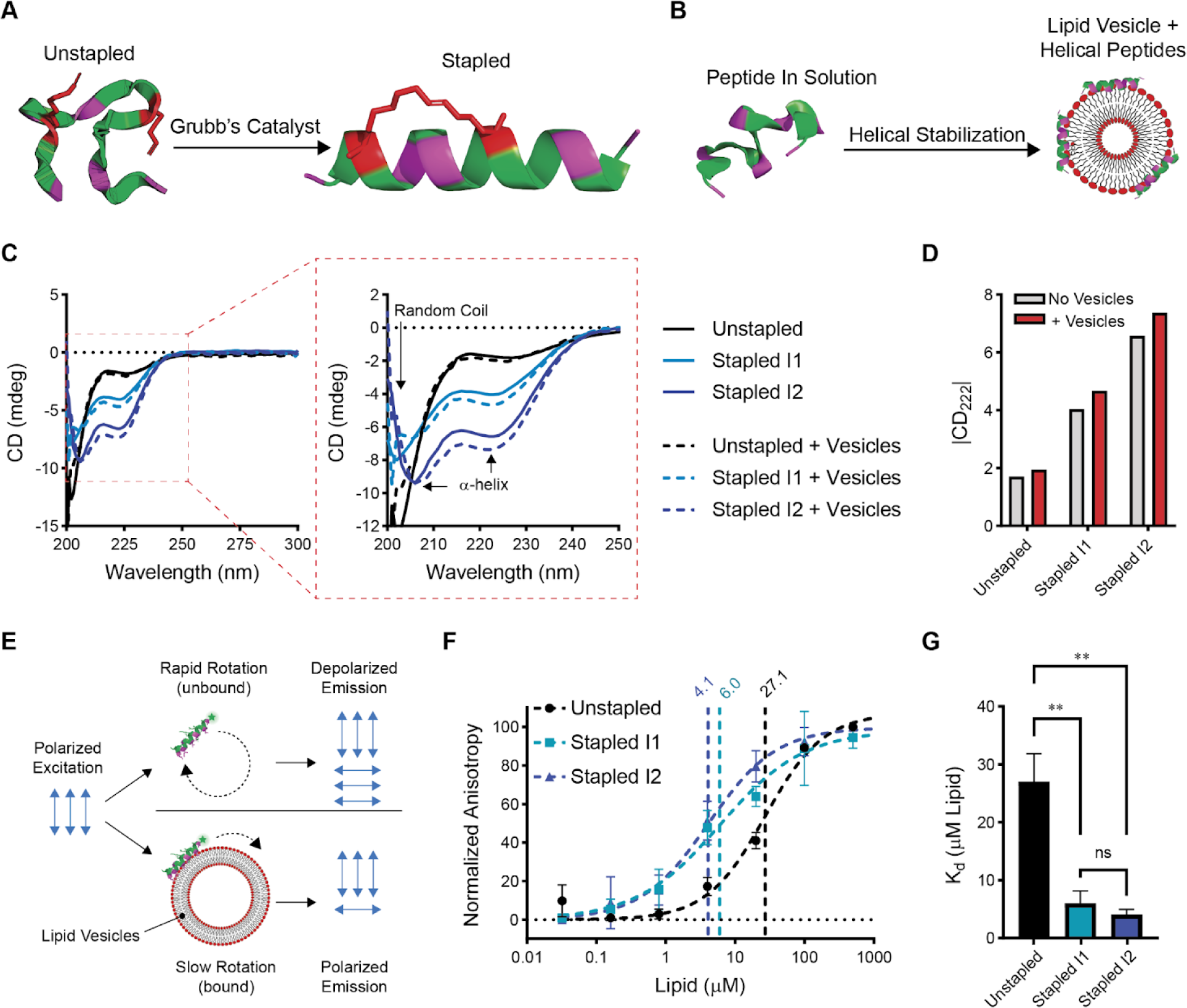
Synthetically Tuning Peptide Helical Content. **(A)** Schematic of synthetic hydrocarbon staple introduced to the peptide to force a helical conformation. **(B)** Schematic of suggested helical stabilization (induced fit) when a peptide binds a lipid membrane. **(C)** Circular dichroism of peptides before and after the addition of lipid vesicles in 50 mM KCl (n=1). **(D)** Quantification of CD at 222 nm showing the increased helical content of the stapled peptides and stabilization after vesicle binding. **(E)** Schematic of fluorescence anisotropy experiment to measure peptide binding to vesicles. **(F)** Fluorescence anisotropy curves of peptides (25 nM) binding to lipid vesicles (n=3). **(G)** Comparison of K_d_ values obtained from anisotropy.

To assess helical content both in solution and after interacting with a membrane we performed circular dichroism of all peptides both with and without 100 nm vesicles (Fig. 4B, C). As described earlier, the unstapled peptide exhibits strong random coil signal on its own as indicated by the sharp negative peak at 200 nm and is mostly unaffected by the addition of vesicles (43). This suggests that the interactions of AH with the membrane are likely minimal in these conditions as we would expect a helical signal if binding occurred (consistent with the simulation results). Interestingly, both stapled isomers exhibited differing degrees of helical signal (minima at 208 and 222 nm) which was then further stabilized after exposure to vesicles. This suggests that while a degree of helical content is needed to be able to bind, the binding event can further stabilize the helix.

Since the helical content is further stabilized during binding, we hypothesize that the binding mechanism can occur partly through an induced fit mechanism i.e., if the peptide is in a partially folded conformation the binding process can induce the conformation to change to a more idealized helix; however, from the unstapled peptide we know that the binding is not strong enough on its own to fold the peptide if it is not aided by other means such as stapling or presumably the context of surrounding amino acids in the native sequence. These observations are in agreement with our FES simulations, discussed earlier; see Figure 3. This change in helical content was assessed through the magnitude of the 222 nm peak (Fig. 4D) showing that the extent of helical stabilization is improved for more helical peptides, likely as a result of higher binding affinity. However, it is important to note that the surfaces of the vesicles are likely saturated with peptides as a high peptide concentration relative to the lipid concentration is needed to obtain clear CD spectra (50μM peptide with 100μM lipid). Excess unbound peptides will reduce the apparent change since not all peptides will be able to adopt the stabilized conformation on the membrane resulting in an underestimate of the true helical stabilization.

To directly measure binding affinity, we fluorescently tagged the peptides using an amine-reactive fluorophore and measured fluorescence polarization l allowing us to detect binding to lipid vesicles (Fig. 4E). Measuring the polarization using 100 nm lipid vesicles with varying lipid concentrations (Fig. S8) allowed us to calculate Kd values for each peptide (Fig. 4F, G), revealing that the stapled peptides have a significantly higher affinity for the membranes. While the difference in Kd between the two stapled isomers is not statistically significant, it is important to note that the trend follows that of helical content. Together these experimental and computational results highlight the importance of helical content for enabling peptide-membrane engagement.

### Including Adjacent Amino Acids Promotes the Folding of the Amphipathic Helix and Membrane Affinity

From the experimental and computational work thus far, it is apparent that the terminal charges and helical folding could have a large impact on the energy barriers for AH folding and ultimately AH binding to the membrane. To examine this in the context of the native protein we explored the addition of 8 amino acids both upstream and downstream of the AH domain (Fig. 5A, S9). We identified that the N-terminal domain has a net neutral charge whereas the C-terminal domain has a net charge of +3. Interestingly when comparing the C-terminal region of the AH domain in several other organisms, it was revealed that the net positive charge on the flanking C-terminal amino acids is a conserved property, which may facilitate binding to the membrane (Fig. S10) (46). In addition to the possibility of the flanking amino acid regions and their charges being involved in binding, these extended domains can have a capping effect and promote helical content (47,48). We tested the synthetic version of this extended peptide (termed 8-AH-8) again using CD (Fig. 5B) and fluorescence polarization (Fig. 5C) to compare to the individual AH domain. The circular dichroism showed a somewhat stronger helical signal compared to the isolated AH domain, but also retained a strong random coil peak suggesting a mixture of folded and unfolded domains. However, despite the random coil peak apparent on CD, the 8-AH-8 peptide exhibits a higher binding affinity for membranes, comparable to that of the stapled AH domains. We speculate that this increased affinity of the 8-AH-8 peptide is due to its core membrane-binding domain (AH) adopting a helical conformation, mediated by the terminal unfolded 8 residues that are exerting a helix-capping effect on the AH domain.

**Figure 5.**
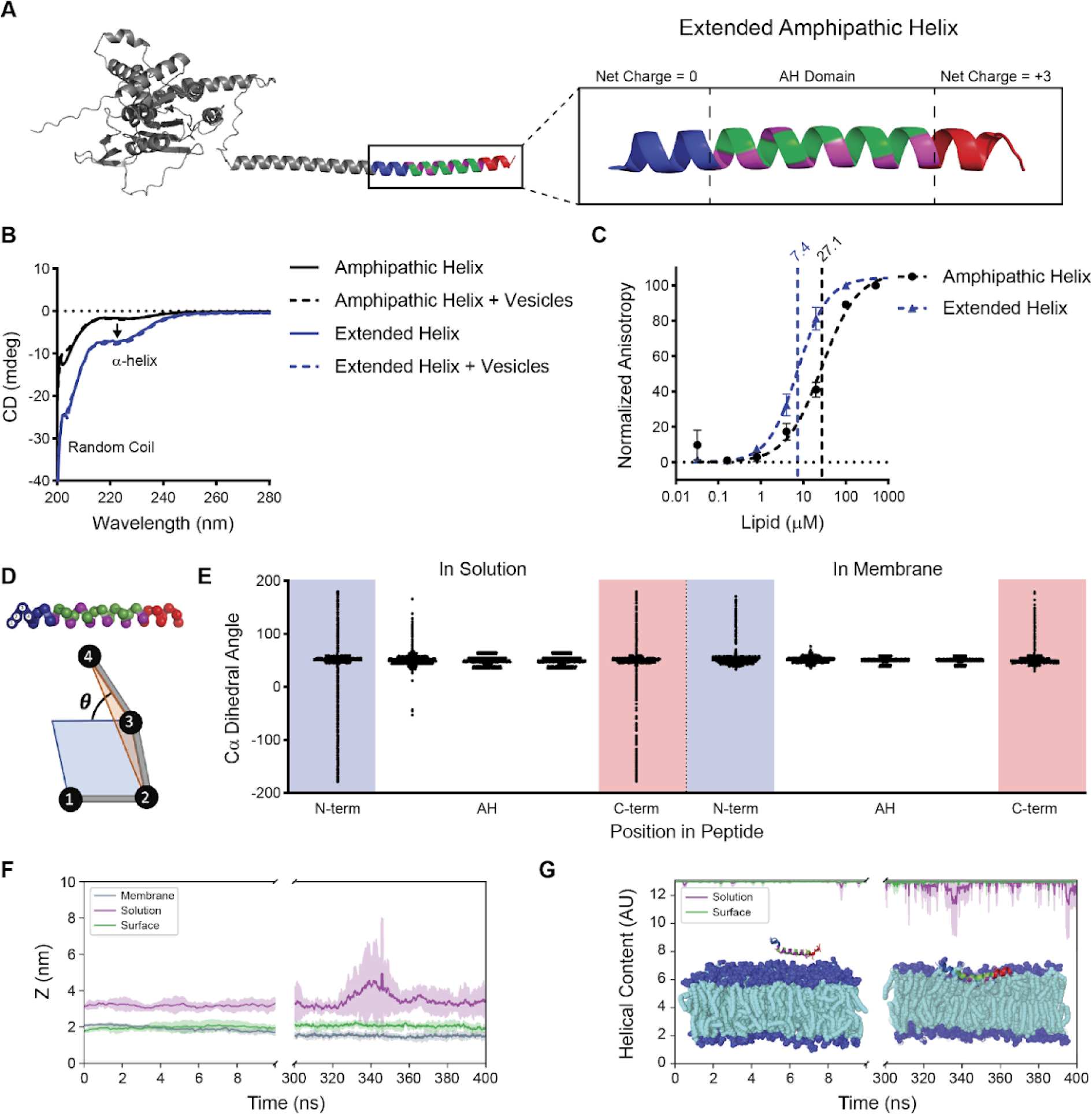
Synthetically Tuning Peptide helical content. **(A)** AlphaFold model of Cdc12 with inset showing the amphipathic helix domain with the flanking 8 amino acids on each side. **(B)** Circular dichroism of peptides before and after the addition of lipid vesicles in 50 mM KCl (n=1). **(C)** Fluorescence anisotropy curves of peptides (25 nM) binding to lipid vesicles (n=3). **(D)** Schematic of how dihedral angles are measured from the peptide backbone over a range of four residues. **(E)** Dihedral analysis of the extended peptide in solution and bound to the lipid membrane (n=4). **(F)** Z-distance calculations from MD simulations showing the relative position of the peptide to the membrane. **(G)** Helical content calculated from MD simulations. For MD analysis, gray represents the membrane, purple represents the peptide initialized in the solution, and green represents the peptide initialized in the membrane.

To test the hypothesis that the N- and C-terminal domains could be flexible while the AH domain remains folded and stable, we performed molecular dynamics simulations of the extended AH initiated in the solution (unbound), and on the surface of the bilayer (bound). To look closer at the flexibility and helical content of the AH, C-, and N-terminal domains, we measured the dihedral angle between each four alpha carbons on the peptide backbone within 34 residues (Fig. 5D), and we averaged the calculated dihedral angle across all four simulation replicas (the average of 4×400 nanoseconds). The dihedral analysis for the extended AH floating in solution shows that the central AH domain remains helical in all four simulation replicas with an angle of 50° (Fig 5E) which indicates a helical secondary structure, based on a survey of dihedral angle analysis of different protein structures from the PDB (49,50). For the N-terminal domain, we observe a wide range of fluctuation of the dihedral angle showing that the N-terminus is very flexible and unfolds in the solution. For the C-terminal domain, on the other hand, the fluctuations are observed only for the residues located at the very end suggesting that the C-terminal domain is more folded and less flexible than the N-terminal domain. This result suggests that the N-terminal domain may be responsible for the random coil signals observed on CD. The dihedral analysis of the simulations initiated from bound extended AH on the bilayer surface mostly shows a 50° dihedral angle throughout the whole peptide residues (Fig. 5E) with only the very ends being flexible. This analysis further supports our suggested induced-fit mechanism where semi-helical peptides in solution are further folded during the binding process.

To further probe the effect of adding upstream and downstream amino acids, we compared the helical content and z-position of the AH domain in the center of extended residues with the isolated AH domain (18-residue sequence: see Fig. 1). The distance of the AH domain center of mass in the extended sequence from the bilayer can be a proxy for the affinity. Comparing the average distance of the AH domain in the extended peptide (Fig. 5F) with the result from the simulations of the AH domain (Fig. 2C) shows that in the solution, the extended peptide remains close to the membrane, while the isolated AH domain escapes away. The extended peptide shows short-lived interactions with the bilayer but does not insert into the membrane in the simulation time span. Looking at the helical content of the AH domain from the extended peptide (Fig. 5G) we observed a significant difference compared with the helical content of the isolated AH domain (Fig. 2D). As indicated by the dihedral analysis, the presence of extended residues helps the AH domain to preserve the helical content in the solution making the peptide more likely to bind to the lipid membrane. As expected, when the peptide is initialized in the membrane (Fig. 5F, G), the peptide remains stably bound with a high degree of helical content.

## DISCUSSION & CONCLUSION

Our combined computational and experimental work has provided a key basis for understanding native septin-membrane interactions. While we have clearly demonstrated the importance of the helical content and charges immediately surrounding the AH domain, it remains to be uncovered how differences in surrounding sequences (length, charge, hydrophobicity) across organisms could impact how each septin behaves and elicits specific membrane curvature preferences. For example, would a septin with a more charged C-terminal sequence have a higher affinity for membranes? Changes to pH or salt concentrations that could either alter or shield charges should be explored in more depth. Additionally, the parameter space to be explored in a biological context is vast. In addition to changes in affinity due to charge of the peptide, the target subcellular membrane composition is likely to have a large impact on how amphipathic helices are incorporated into the bilayer. The DOPC:PLPI lipid composition used here and in previous studies is greatly simplified compared to a cell membrane where the rich variety of lipid headgroups and membrane mechanical properties it exhibits depends on time and location. Changing lipid composition will alter the size distribution, density and lifetime of lipid packing defects, where a higher density of larger and longer-lived defects is expected to increase AH incorporation and time bound to the membrane (10,51).

Unfolded AH domains in solution likely go through a stepwise process of folding while inserting into the membrane where lipid binding induces helix formation, which in turn promotes more favorable contacts with membrane lipids (4,52). This transition from unstructured in solution to alpha helical in the membrane is shared by non-curvature sensing peptides as well (53). This is in good agreement with simulation data suggesting lipid packing defects are too small to allow for the entire hydrophobic face of an AH domain to embed into the membrane but are more likely to accommodate a single hydrophobic side chain at a time (4). While it is possible for membranes to mediate folding of otherwise unstructured protein domains, our results here for septins suggest that amino acids surrounding their AH domain promote helical folding, even in solution, and stapling the peptide to restrict it into helical conformation can actually increase its affinity for the membrane. Work on the only other micron-scale curvature sensing AH domain in SpoVM proteins also shows an increase in membrane affinity when the helix is stabilized through a mutation (9). Combined, this supports a model where septins (and potentially other micron-scale curvature-sensing domains) may exhibit a partially folded AH domain in the context of the protein complex and independent of membrane binding. In this case, rather than using an iterative process of lipid binding and folding, septins may rely on the rearrangement of membrane lipids upon contact of a folded AH, which in turn makes modest contributions to the helical content of the AH domain, in agreement with the small changes observed by CD. While these membrane defects may not be large enough to initially insert the entire hydrophobic face of the helix into the membrane, it is possible the folded helices also undergo a stepwise binding process where a portion of the hydrophobic face binds to a small defect followed by the formation of additional defects to allow the complete insertion.

Computationally, problems still exist with the size of the required search space for free energy calculations, and the speed at which these calculations can be performed. Enhanced sampling methods allow us to find FES without needing to solve the ‘forward problem’ of watching peptides bind to the membrane. However, these calculations are still time expensive. Multi-walker metadynamics simulations were carried out using dual-socket Intel Icelake processors, and still required >1 month of real-world runtime to converge. This was with 4 independent walkers using 10 nodes (1,840 cores). This makes running multiple charge-cap choices, amino acid sequence, salt concentration, etc, prohibitively expensive to simulate.

Expanding from a single AH domain into investigating a tandem series of multivalent AH and its effect on membrane binding will be key to understanding septin’s curvature sensing/inducing properties. Should simulations enable us to observe how the multivalent display of AH domains will influence membrane interactions, we could tune proteins or synthetic materials to have various curvature sensing and inducing characteristics. For example, AH-anchored on DNA origami (54) could be used to mimic septin octameric and filamentous states, with the ability to tune the density (spacing) of AH domains. This type of structure might exhibit tunable affinities for preferred curvatures and enable unprecedented regulation of living or synthetic cell shapes.

## AUTHOR CONTRIBUTIONS

C.J.E. performed simulations and enhanced sampling of AH domains, S.J.K. performed experimental characterization of structure and vesicle binding of all peptides. S.M.M. performed simulations and enhanced sampling of AH domains and Extended AH domains. Q.W. assisted with peptide synthesis and purification. S.B. performed dynamic light scattering of vesicles. E.J.D.V., B.N.C, W.S, S.M.H., M.G.F., D.K., and A.G. provided helpful input and discussion. R.F. and E.N. supervised the work.

## DECLARATION OF INTERESTS

The authors declare no conflicting interests.

## ACKNOWLEDGEMENTS

Dynamic light scattering, circular dichroism, and fluorescence anisotropy were performed at UNC’s Macromolecular Interactions Facility supported by the National Cancer Institute of the National Institutes of Health under award number P30CA016086. Simulations in this paper were carried out at the Flatiron Institute computing clusters. The content is solely the responsibility of the authors and does not necessarily represent the official views of the National Institutes of Health. We thank the University of North Carolina’s Department of Chemistry Mass Spectrometry Core Laboratory which is supported in part with funding from the University of North Carolina’s School of Medicine Office of Research. This work was supported by the Alfred P. Sloan Foundation Grant G-2021-14197. We would like to thank Pilar Cossio, and Abhilash Sahoo for their useful input in discussions.

## SUPPLEMENTAL MATERIAL

**Figure S1.**
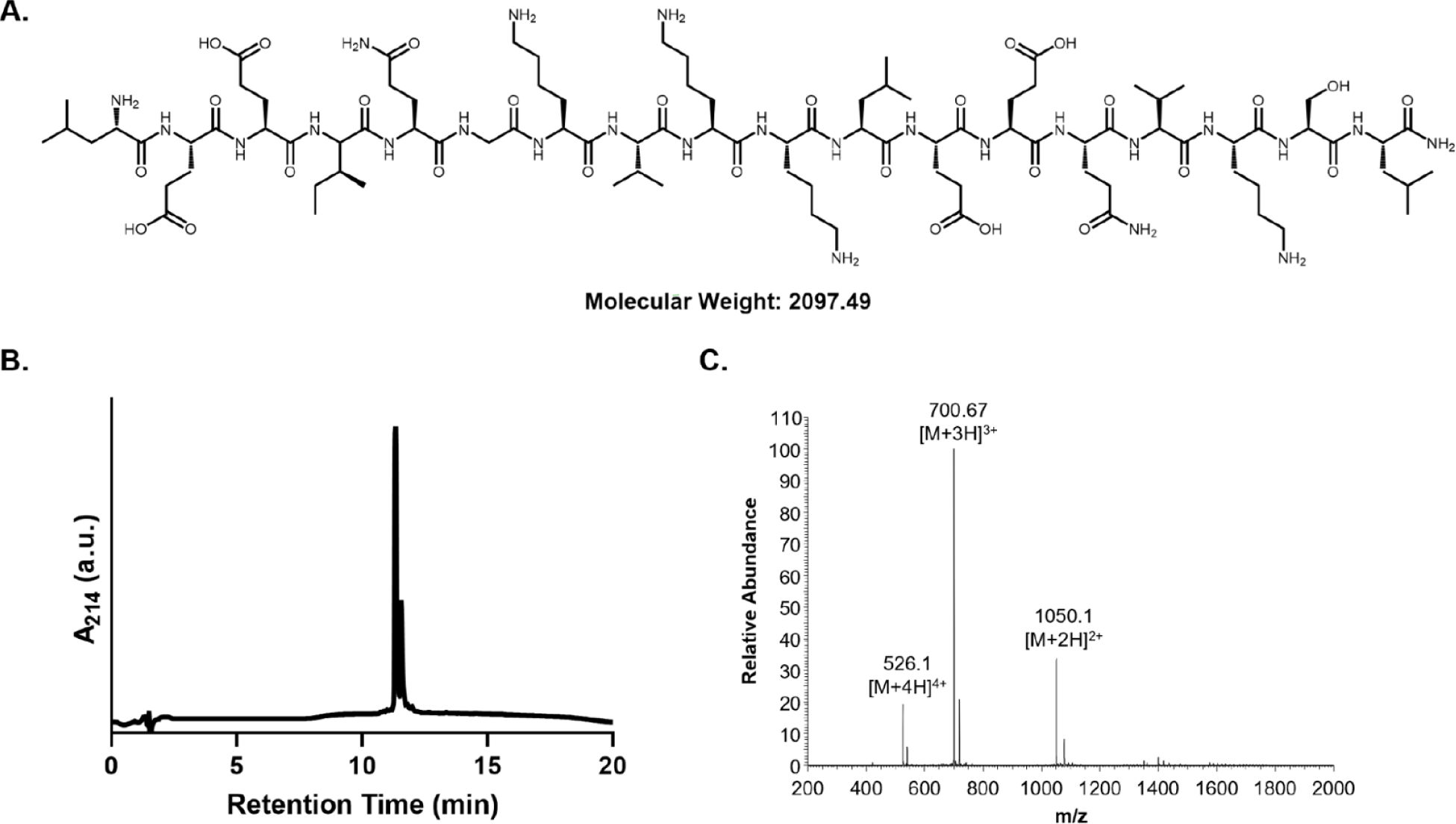
Synthesis and Purification of AH Monomer. Chemical structure (A) of AH monomer. Analytical HPLC trace (B) showing peptide purity. Mass spectrum (C) of AH peptide confirming identity.

**Figure S2.**
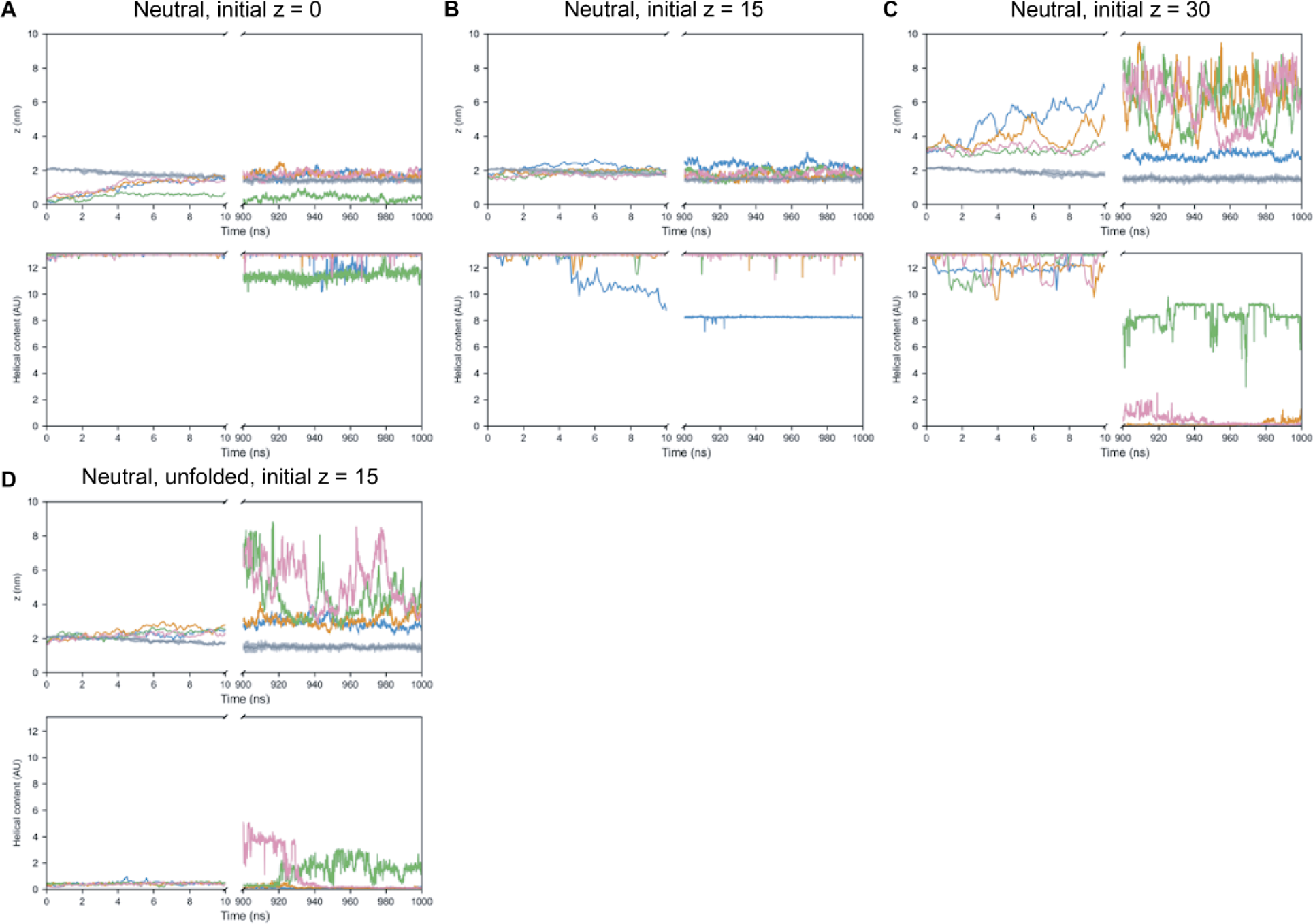
Simulations of neutral-capped amphipathic helix in different starting configurations demonstrate stable equilibrium at the surface of the membrane. Top row: individual run center-of-mass to helix center-of-mass distance as a function of time. Bottom row: individual run helical content as a function of time. Colored lines show the individual simulations runs. Gray curve in top row shows the average location of the membrane phosphate groups with the shaded regions showing the standard error. (A) Folded amphipathic helices placed in the middle of the membrane (center-of-mass distance 0 nm away from the center of the membrane) go to either the surface of the membrane or wind up tilted acting as pseudo-trans-membrane proteins (green curve). However, none unbind during the course of the simulations, and all remain helical. (B) Folded amphipathic helices placed near the surface of the membrane (center-of-mass distance 1.5 nm away from the center of the membrane) remain bound to the surface and remain mostly helical (blue curve being the exception). (C) Folded amphipathic helices placed in solution (center-of-mass distance 3 nm away from the center of the membrane) unfold over time and do not bind to the membrane. (D) Unfolded amphipathic helices placed near the surface of the membrane (center-of-mass distance 1.5 nm away from the center of the membrane) are ejected from the membrane and do not fold into an α-helix configuration. (N=4) neutral-cap simulations were run for 1 μs with 50 mM KCl at 300K.

**Figure S3.**
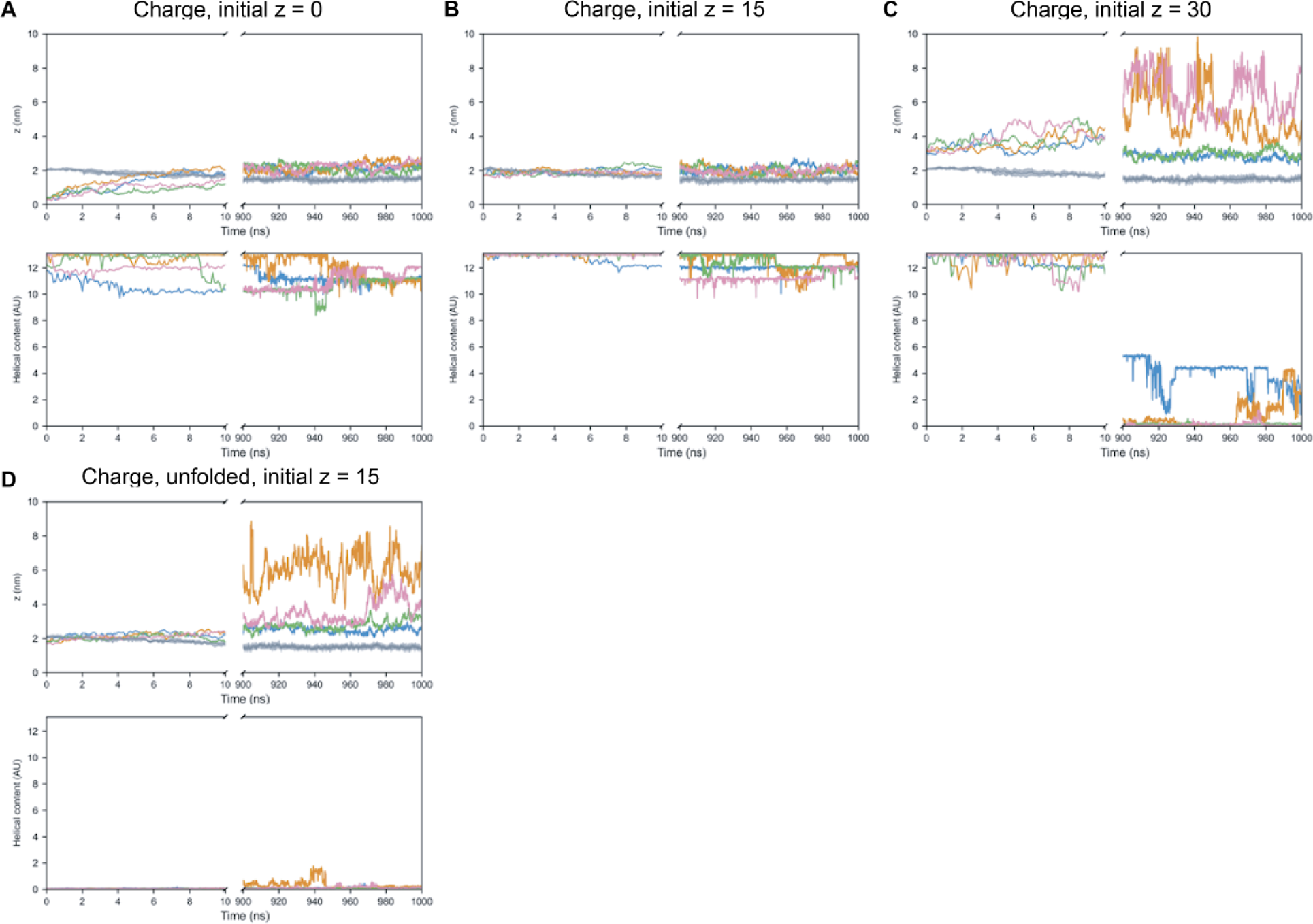
Simulations of charge-capped amphipathic helix in different starting configurations demonstrate stable equilibrium at the surface of the membrane. Top row: individual run center-of-mass to helix center-of-mass distance as a function of time. Bottom row: individual run helical content as a function of time. Colored lines show the individual simulations runs. Gray curve in the top row shows the average location of the membrane phosphate groups with the shaded regions showing the standard error. (A) Folded amphipathic helices placed in the middle of the membrane (center-of-mass distance 0 nm away from the center of the membrane) go to the surface of the membrane and do not unbind or unfold during the course of the simulation. (B) Folded amphipathic helices placed near the surface of the membrane (center-of-mass distance 1.5 nm away from the center of the membrane) remain bound to the surface and remain helical during the course of the simulation. (C) Folded amphipathic helices placed in solution (center-of-mass distance 3 nm away from the center of the membrane) unfold over time and do not bind to the membrane. (D) Unfolded amphipathic helices placed near the surface of the membrane (center-of-mass distance 1.5 nm away from the center of the membrane) are ejected from the membrane and do not fold into an α-helix configuration. (N=4) charge-cap simulations were run for 1 μs with 50 mM KCl at 300K.

**Figure S4.**
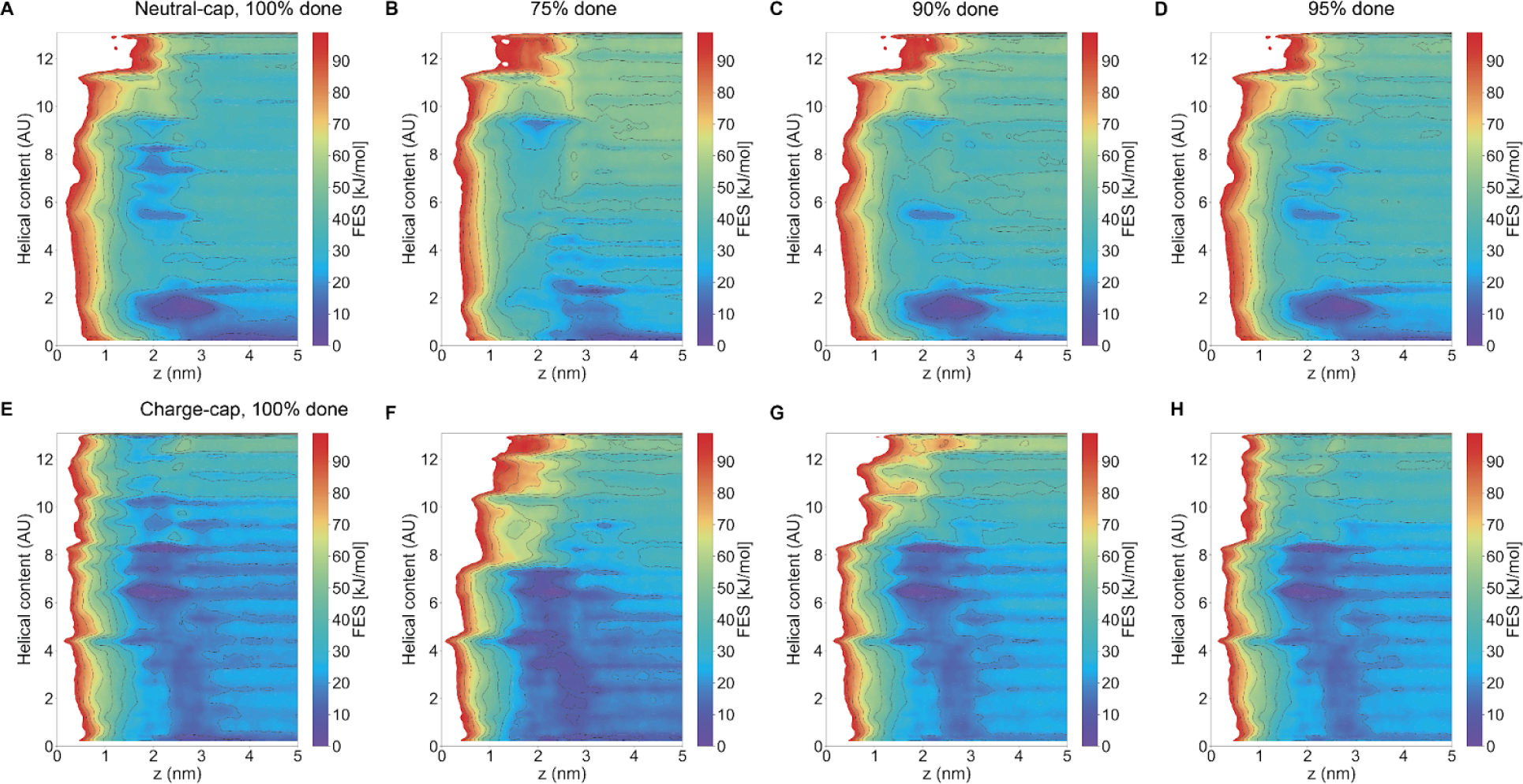
Convergence testing for metadynamics based on block averaging at different timepoints during the metadynamics simulation. (A-D) Neutral-capped peptide convergence tests. (E-H) Charge-capped peptide convergence tests. (A,E) Reconstructed FES from gaussian hills deposited during metadynamics algorithm, same as Figure 3A,B. (B,F) Metadynamics runs at 75% complete. (C,G) Metadynamics runs at 90% complete. (D,H) Metadynamics runs at 95% complete. The 90% and 95% complete runs show convergence to near the final FES.

**Figure S5.**
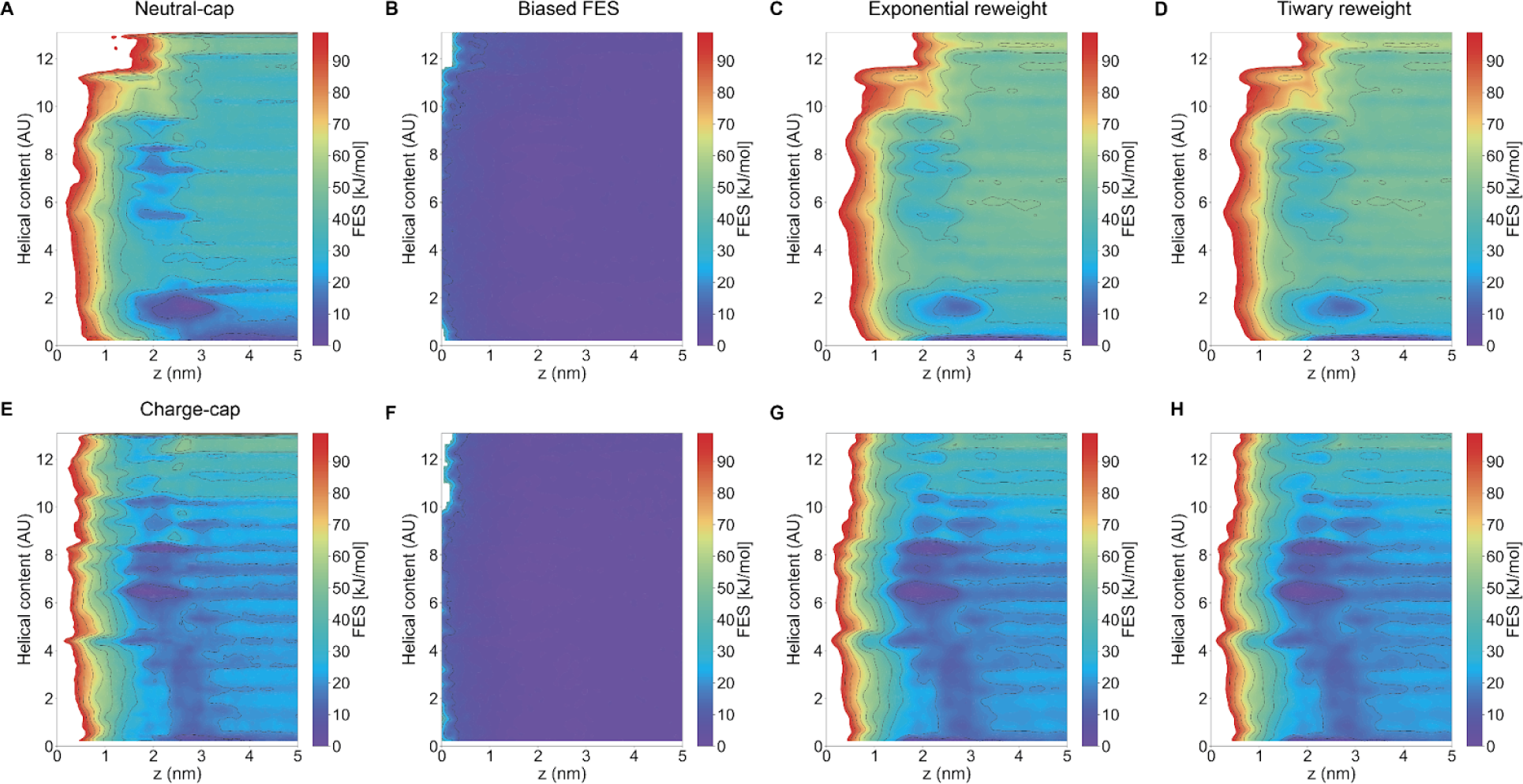
Convergence testing for metadynamics based on biased free energy surface and reweighting procedures. (A-D) Neutral-capped peptide convergence tests. (E-H) Charge-capped peptide convergence tests. (A,E) Reconstructed FES from gaussian hills deposited during metadynamics algorithm, same as Figure 3A, B. (B, F) Biased FES sampling the probability density of different states during metadynamics simulation. If metadynamics has converged, this should be flat. For the regions of interest, this shows that metadynamics has converged up to an error of around 10 kJ/mol for both neutral- and charge-capped AH-domains. (C, G) FES reconstructed via an exponential reweighting procedure in PLUMED. (D, H) FES reconstructed via Tiwary reweighting procedure in PLUMED. Exponential and Tiwary reweighting procedures show a similar result to the FES constructed by summing up the gaussian hills.

**Figure S6:**
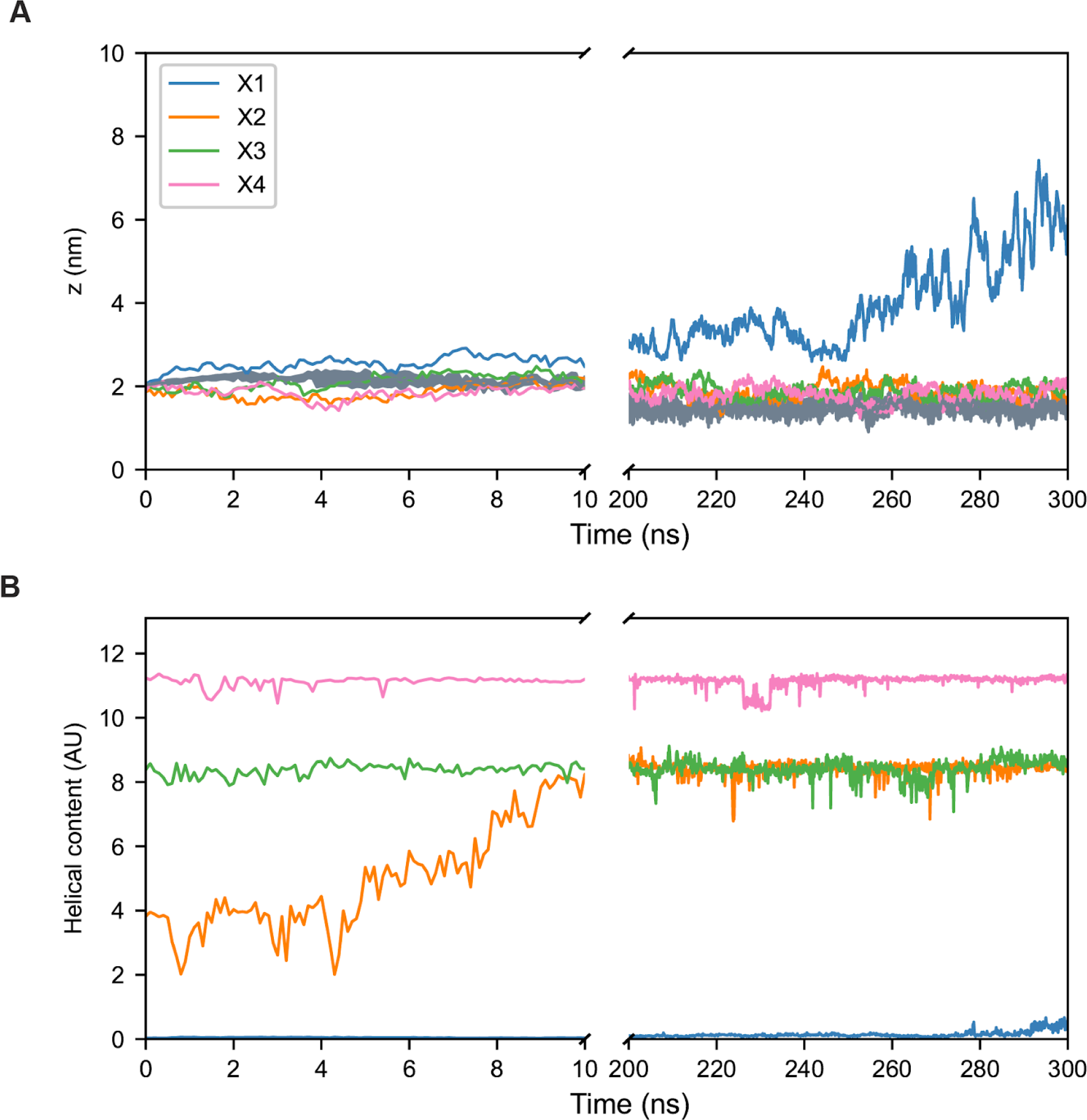
MD simulations of points on free energy surfaces. Validating trajectories of simulations on the free energy surface. Four points (X1, (z ∼ 1.9 nm, alpha ∼ 0), blue; X2, (z ∼ 1.9 nm, helical content ∼ 4), orange; X3, (z ∼ 1.9 nm, helical content ∼ 8), green; and X4, (z ∼ 1.9 nm, helical content ∼ 12), pink)) were chosen on the FES corresponding to neutral-capped AH domains. (A) Z-position of AH domain. AH domains with no helical content are ejected from the membrane during the simulation (X1), while the others remain bound (X2-X4). (B) Helical content of AH domain. AH with no initial helical content remains non-helical (X1), while a lower helical content AH acquires more helical content over the simulation (X2).

**Figure S7.**
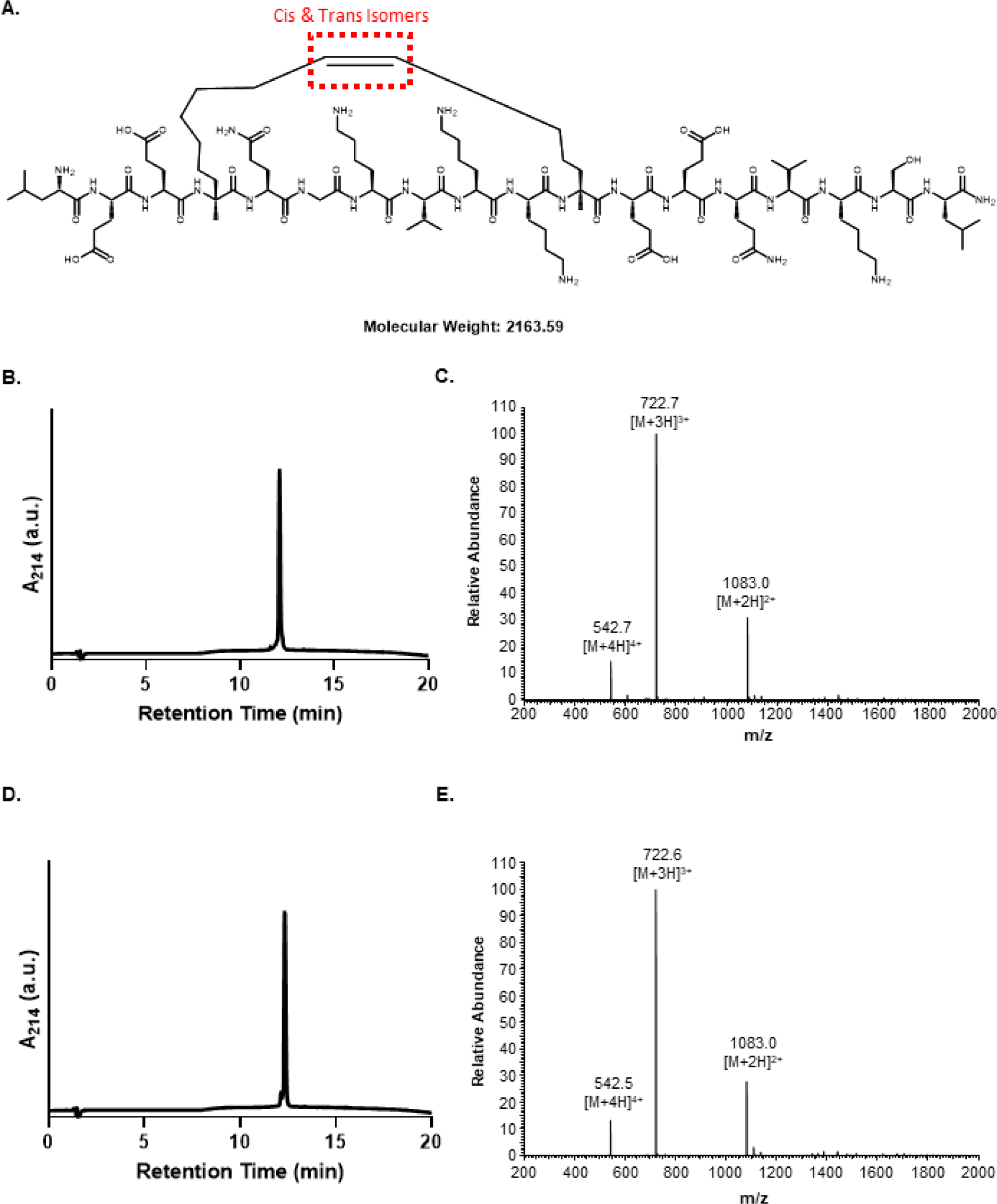
Synthesis and Purification of Stapled AH Peptide. Chemical structure (A) of stapled AH peptide showing the location of the double bond which yields cis-trans isomers. Analytical HPLC trace (B) of isomer 1 showing purity and mass spectrum (C) confirming identity. Analytical HPLC trace (D) of isomer 2 showing purity and mass spectrum (E) confirming identity.

**Figure S8.**
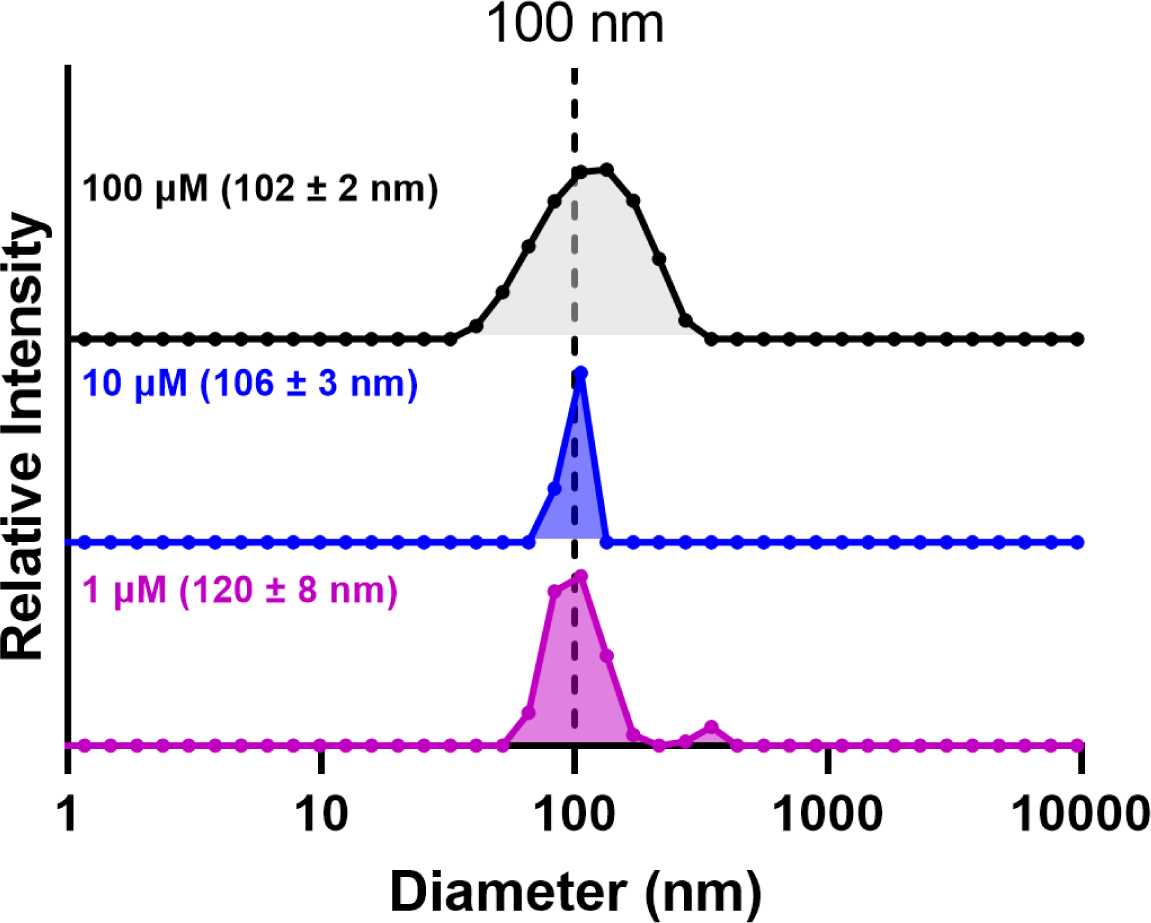
Dynamic Light Scattering of Vesicles. To determine the size distribution of vesicles, dynamic light scattering was performed on vesicles at 1, 10 and 100 µM.

**Figure S9.**
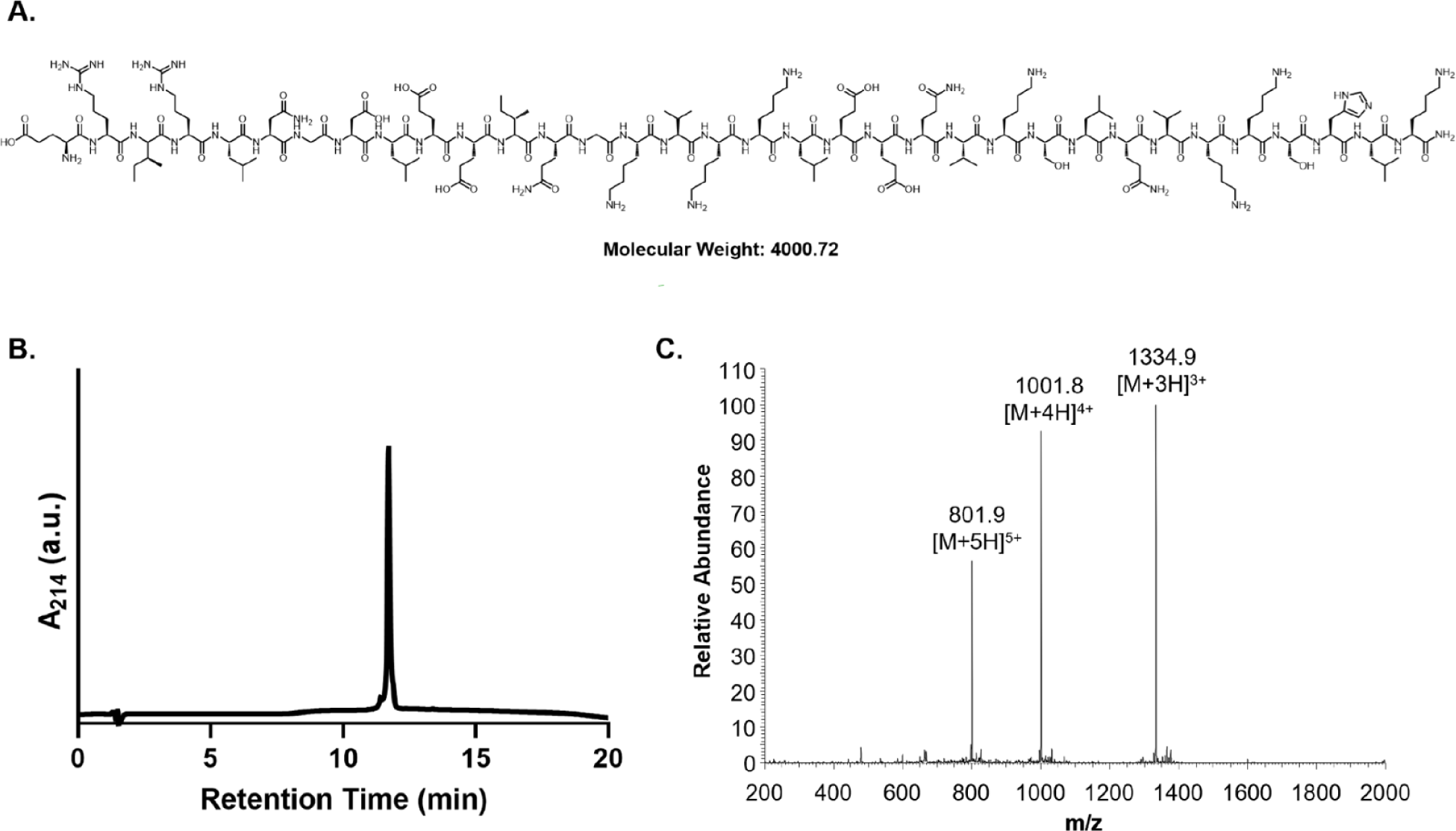
Synthesis and Purification of Extended AH peptide (8-AH-8). Chemical structure (A) of extended AH peptide. Analytical HPLC trace (B) showing peptide purity. Mass spectrum (C) of 8-AH-8 peptide confirming identity.

**Figure S10.**
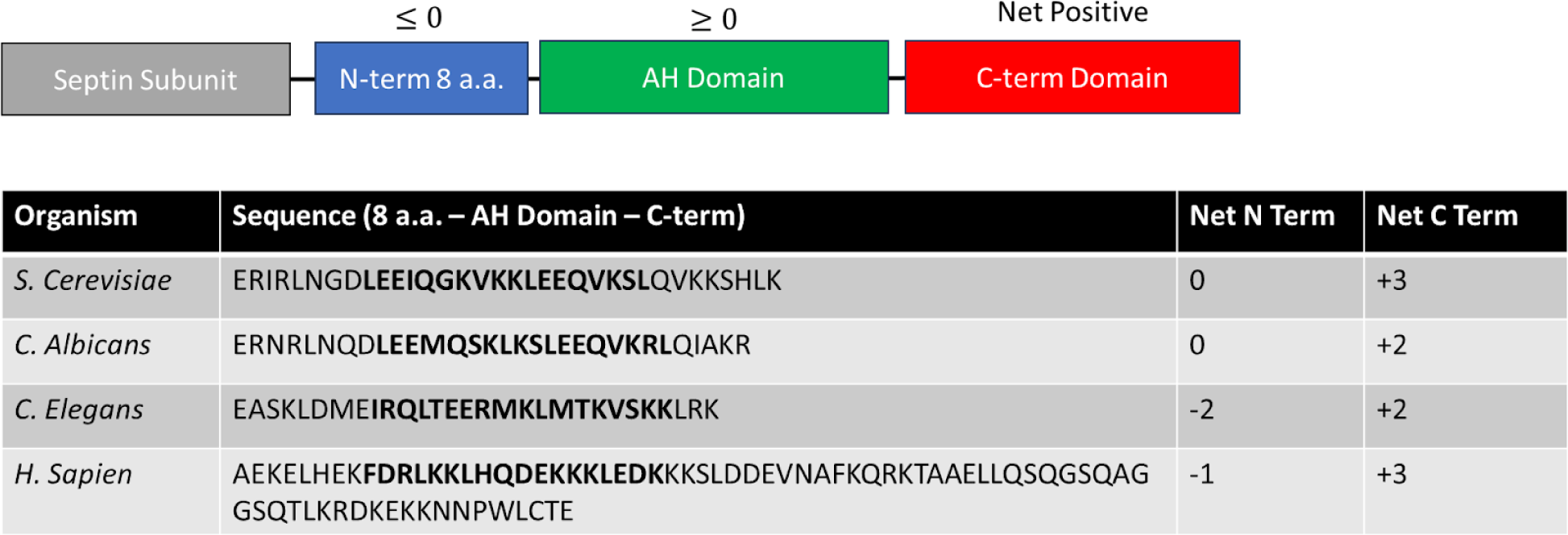
Comparison of septin charges. Comparing the regions surrounding the amphipathic helix domains in septins across organisms revealed that all C-terminal domains have a net positive charge. Sequences were retrieved from Uniprot.

**Table S1.**
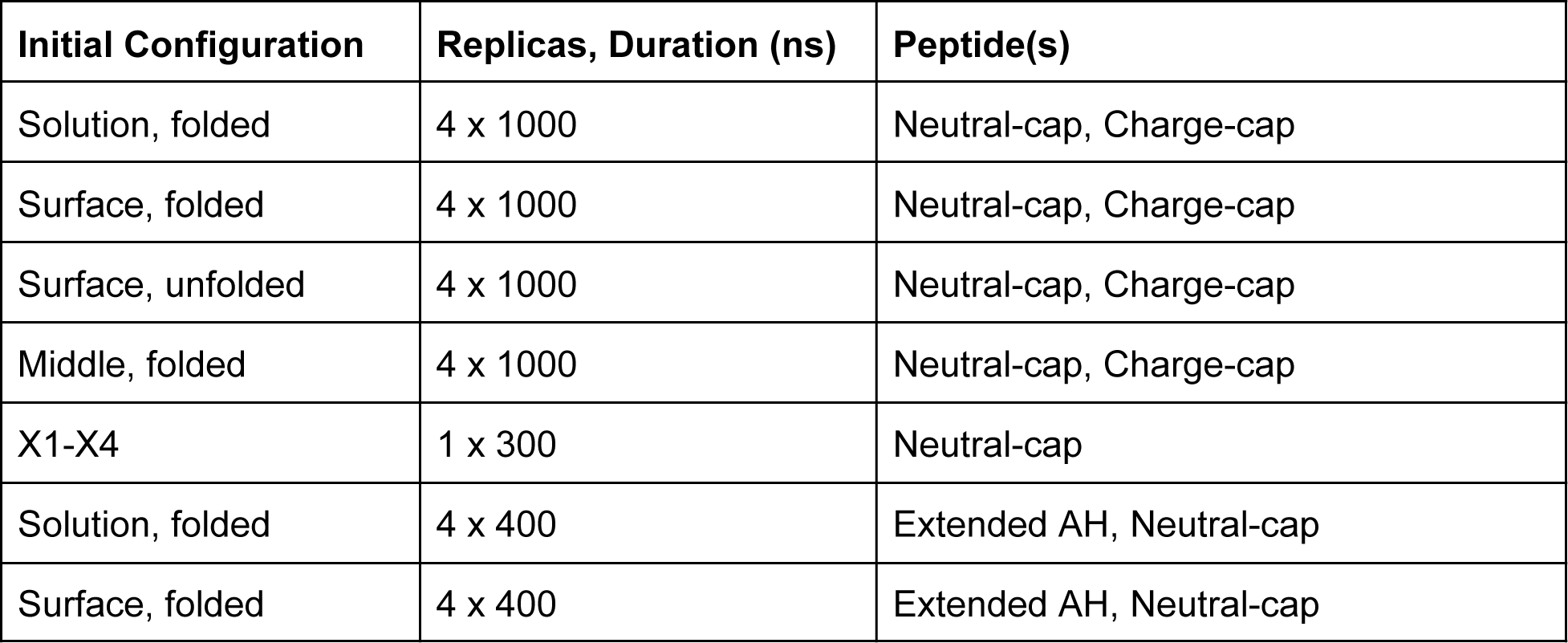
Summary of Molecular Dynamics Simulations.

| Initial Configuration | Replicas, Duration (ns) | Peptide(s) |
| --- | --- | --- |
| Solution, folded | 4 x 1000 | Neutral-cap, Charge-cap |
| Surface, folded | 4 x 1000 | Neutral-cap, Charge-cap |
| Surface, unfolded | 4 x 1000 | Neutral-cap, Charge-cap |
| Middle, folded | 4 x 1000 | Neutral-cap, Charge-cap |
| X1-X4 | 1 x 300 | Neutral-cap |
| Solution, folded | 4 x 400 | Extended AH, Neutral-cap |
| Surface, folded | 4 x 400 | Extended AH, Neutral-cap |

**Table S2.** Summary of Metadynamics Simulations.

| Walkers, duration (ns) | Peptide(s) |
| --- | --- |
| 4 x 3000 | Neutral-capped |
| 4 x 3000 | Charge-capped |

## Notes

### Competing Interest Statement

The authors have declared no competing interest.

